## SupplementaryInformation for "Neural attentional-filter mechanisms of listening success in middle-aged and older individuals"

### for

### Supplementary Notes

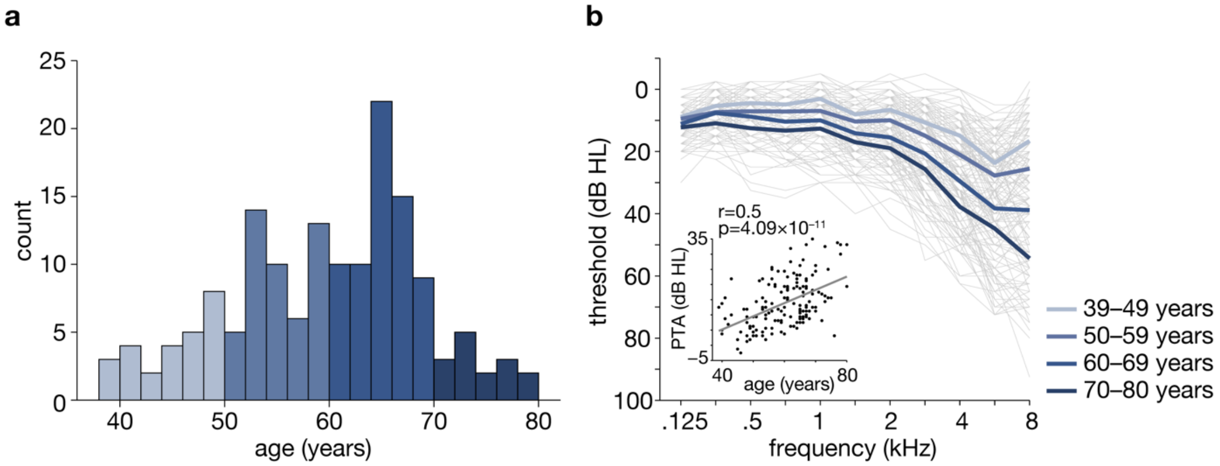


**Supplementary Fig 1. Age and hearing loss distribution.**
**(a)** Histogram showing age distribution of N=155 participants across 2-year age bins.
**(b)** Individual and mean air conduction thresholds averaged across the left and right ear. Thin grey lines show air conduction pure-tone thresholds for individual participants, thick coloured lines indicate average thresholds grouped across four age bins. Inset scatterplot shows positive Pearson correlation ($r(153) = .5$, $p = 4.094 \times 10^{-11}$) of pure-tone average (PTA; mean across left and right ear at 500, 1000, 2000 and 4000 Hz) and age. Black dots show thresholds for individual participants. Source data are provided as a Source Data file.

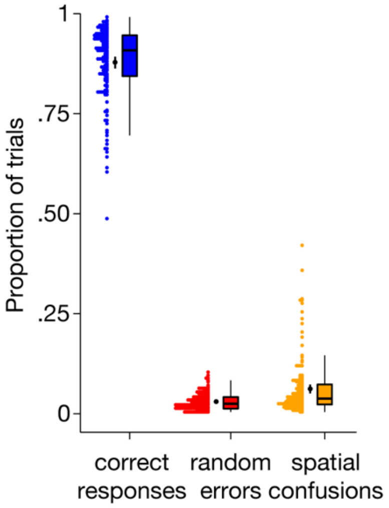


**Supplementary Fig 2. Analysis of response types.** Responses split into correct responses (blue), random errors (red) and spatial stream confusions (orange). Coloured dots are individual proportion of trials for *N*=155 participants, black dots and vertical lines show group means and bootstrapped 95 % confidence interval. Box plots show median centre line, 25th to 75th percentile hinges, whiskers show minimum and maximum within 1.5 × interquartile range. Source data are provided as a Source Data file.

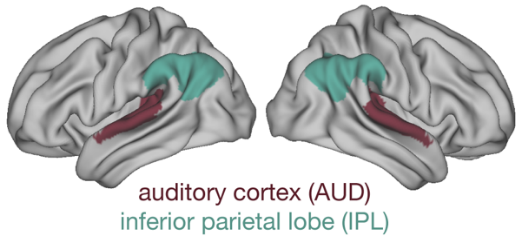


**Supplementary Fig 3. Regions of interest.**
Spatial extent of bilateral region of interest (ROI) in the wider auditory cortex (dark red) and a control ROI in the inferior parietal lobule (green). Source data are provided as a Source Data file.

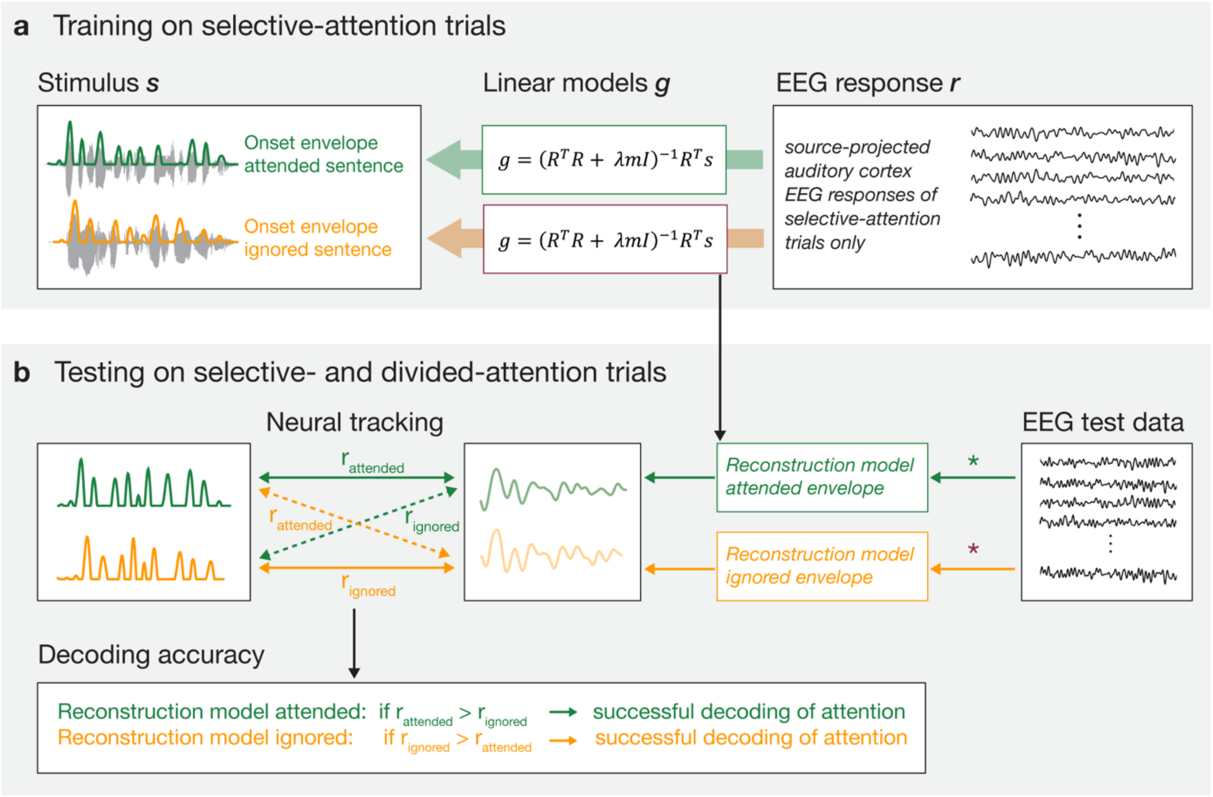


**Supplementary Fig 4. Training and testing of envelope reconstruction models.**
**(a)** Linear backward models were estimated using single-trial onset envelopes and preprocessed EEG responses of selective attention trials, only. Single-subject models for the attended and ignored speech stream were trained separately.
**(b)** Following a leave-one-out procedure for selective attention trials, single-trial envelope reconstructions were computed by convolving the EEG signal of the test trial with the trained reconstructions models averaged across all but the tested trial. For the reconstruction of envelope presented in divided attention trials, we used the average of all single-trial decoder models estimated in training. Neural tracking was quantified as the Pearson-correlation coefficient of the reconstructed and the actually presented envelopes yielding four coefficients per trial and subject. Decoding accuracy was assessed separately for the reconstruction model of the attended and ignored envelope, respectively. For the former, a participant's attention was correctly decoded if the correlation coefficient for the comparison of the reconstructed with the attended envelope ($r_{attended}$) was higher than that of the comparison with the ignored envelope ($r_{ignored}$). The opposite relationship holds for the decoding accuracy of the ignored envelope reconstruction model. Source data are provided as a Source Data file.

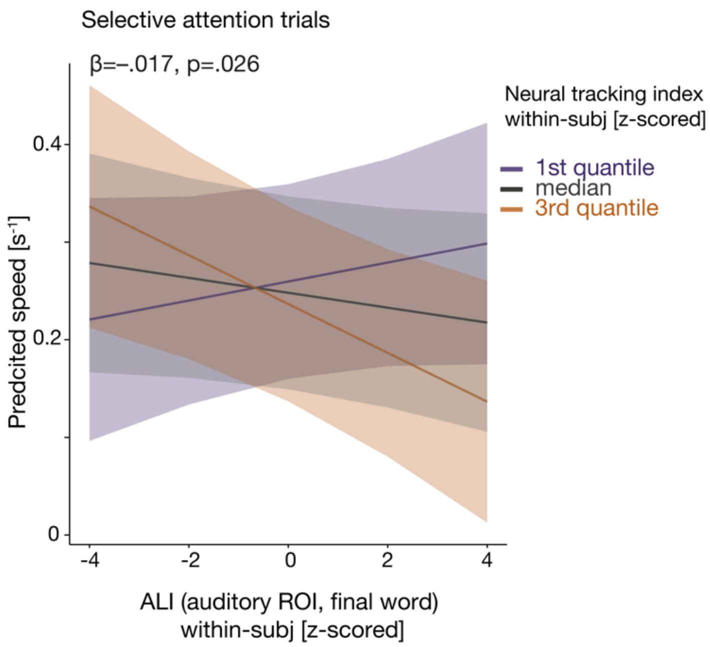


**Supplementary Fig 5. Neural attentional filters predict response speed.**
Under selective attention, the combination of within-subject variability of neural tracking and alpha power lateralization during final word presentation predicts response speed. Predicted effect of alpha power lateralization on response shown for three different levels of neural tracking across all *N*=155 participants. Coloured lines show fixed (group-average) effects, shaded coloured areas indicate 95% confidence intervals.
*β:* slope parameter estimate from general linear mixed-effects model, p-value based on two-sided Wald test, FDR-corrected. Source data are provided as a Source Data file.

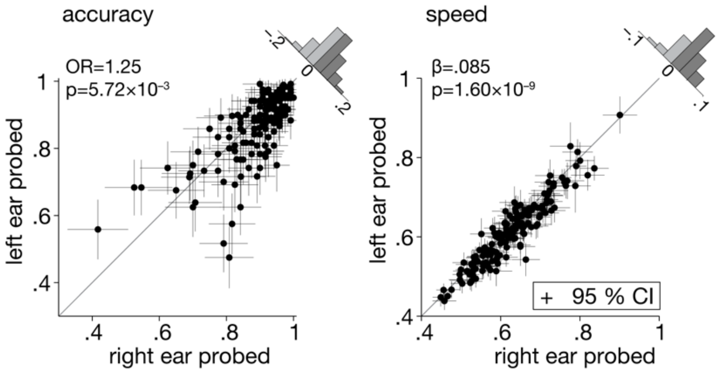


**Supplementary Fig 6. Right-ear advantage for accuracy and response speed.**
Individual right-ear advantage for *N*=155 participants displayed separately for accuracy and response speed, respectively. Black dots indicate individual trial-averages with bootstrapped 95% confidence interval error bars. Histograms show the distribution of the difference of right-ear vs. left-ear probed trials across all participants.
OR: Odds ratio parameter estimate from generalized linear mixed-effects models; *β:* slope parameter estimate from general linear mixed-effects models; p-values based on two-sided Wald test, FDR-corrected. Source data are provided as a Source Data file.

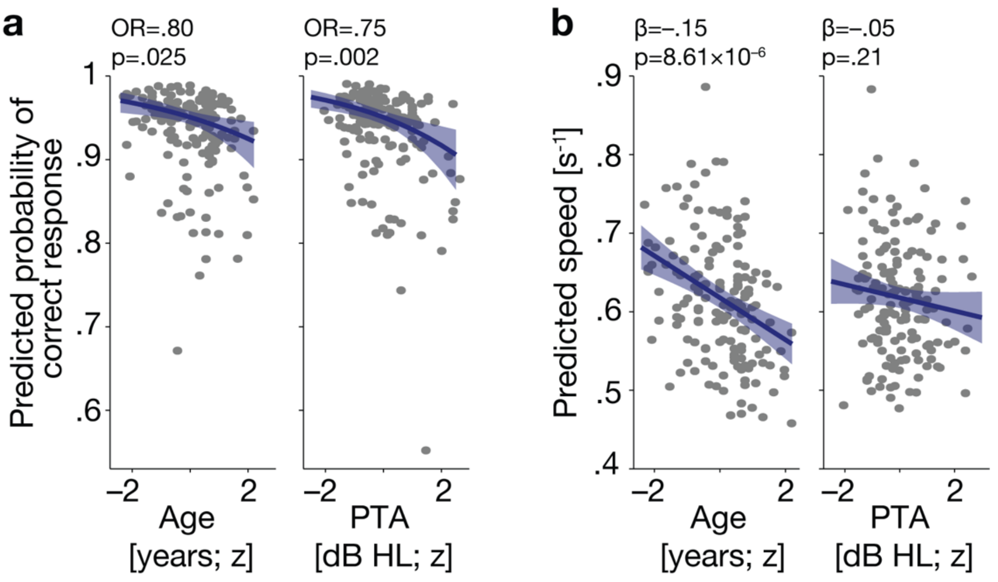


**Supplementary Fig 7. Effects of age and PTA on accuracy and response speed.**
**(a)** Significant effect of age and PTA on single-subject average accuracy. Shown for $N$=155 participants. Grey dots indicate single-subject model predictions for the probability of a correct response. Blue curves reflect fixed (group-average) effects of age and PTA as indicated by the best-fitting generalized linear mixed model (see Table S1 for full model details) along with 95% confidence interval (CI), $p$-values based on two-sided Wald test, FDR-corrected.
**(b)** Significant effect of age on single-subject average on response speed and the absence of a corresponding effect of PTA. Shown for $N$ =155 participants. Grey dots indicate single-subject model predictions for single-subject average speed. Blue curves reflect fixed (group-average) effects of age and PTA as indicated by the best-fitting generalized linear mixed model (see Table S2 for full model details) along with 95% CI, $p$-values based on two-sided Wald test, FDR-corrected. Source data are provided as a Source Data file.

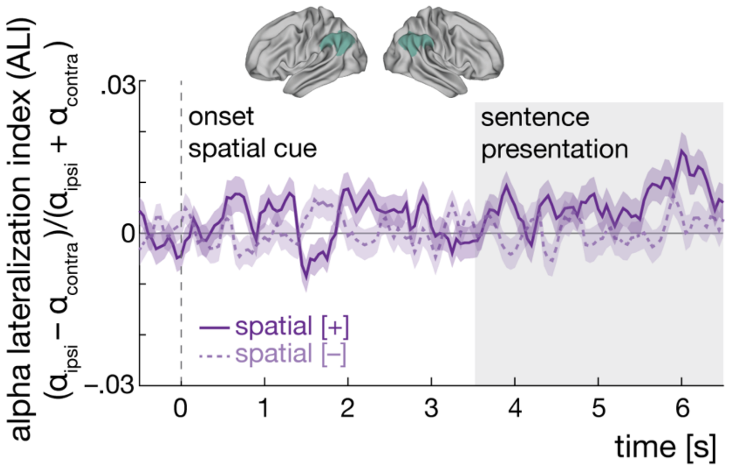


**Supplementary Fig 8. Alpha lateralization in inferior parietal cortex.**
Attentional modulation of 8–12 Hz inferior parietal alpha power throughout the trial for $N$=155 participants. Brain models indicate the spatial extent of the inferior parietal control region of interest (shown in green). Purple traces show the grand-average alpha lateralization index (ALI) for the informative (solid dark purple line) and uninformative spatial cue (dashed light purple line), each collapsed across semantic cue levels. Error bands indicate ± 1 SEM. Positive values indicate relatively higher alpha power in the hemisphere ipsilateral to the attended/-probed sentence compared to the contralateral hemisphere. Shaded grey area shows the time window of sentence presentation. Source data are provided as a Source Data file.

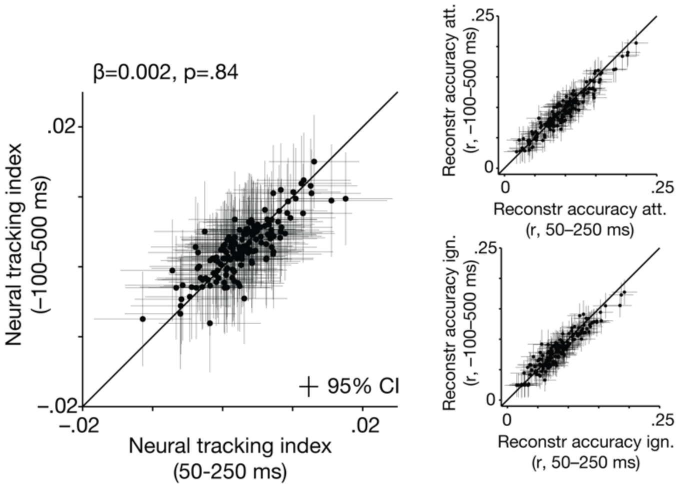


**Supplementary Fig 9. Effect of time lag range on neural tracking strength.** Comparison of single-subject (*N*=155 participants) mean neural tracking as a function of different time lag ranges used for reconstruction. Left panel compares mean neural tracking index per subject (black dots) based on time lags of 50–250 ms vs. −100-500 ms. *β:* slope parameter estimate from general linear mixed-effects model p-value based on Wald test. Error bars reflect 95 % confidence interval. Black dots in right panels show single-subject (*N*=155 participants) mean reconstruction accuracy of the attended envelope (top), and the ignored envelope (bottom) for both ranges of time lags. Error bars reflect 95 % confidence interval. Source data are provided as a Source Data file.

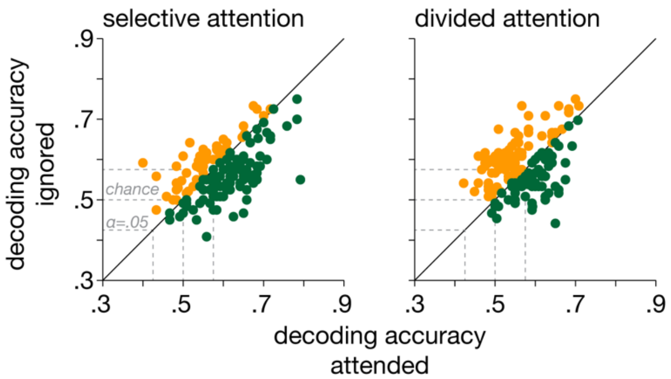


**Supplementary Fig 10. Decoding accuracy per attention condition.** Single-subject (*N*=155) decoding accuracy of the attended and ignored reconstruction models in selective-attention (left panel) and divided-attention trials (right panels). Green dots below the 45° line indicate that for these subjects decoding accuracy for the attended envelope was higher than for the ignored envelope, and vice versa for red dots above the 45° line. Grey dotted lines indicate performance at chance level, and significantly above/below chance level based on a binomial test at α=.05. Source data are provided as a Source Data file.

**Supplementary Table 1: Predicting single-trial accuracy**

| | Accuracy | | | | | Probed-R trials | | | | | Probed-L trials | | | | |
|---|---|---|---|---|---|---|---|---|---|---|---|---|---|---|---|
| *Predictors* | *Odds Ratios* | *std. Error* | *CI* | *z-value* | *p* | *Odds Ratios* | *std. Error* | *CI* | *z-value* | *p* | *Odds Ratios* | *std. Error* | *CI* | *z-value* | *p* |
| Intercept | 19.156 | 0.084 | 16.236 – 22.602 | 34.986 | **8.21e-267** | 22.364 | 0.094 | 18.605 – 26.883 | 33.092 | **7.07e-239** | 16.954 | 0.094 | 14.091 – 20.397 | 30.001 | **1.72e-196** |
| Spatial cue (selective) | 3.398 | 0.117 | 2.702 – 4.273 | 10.466 | **1.36e-24** | 3.593 | 0.141 | 2.725 – 4.738 | 9.063 | **1.15e-18** | 3.071 | 0.126 | 2.399 – 3.932 | 8.904 | **4.85e-18** |
| Semantic cue (specific) | 1.099 | 0.106 | 0.892 – 1.354 | 0.886 | 6.88e-01 | 1.194 | 0.120 | 0.944 – 1.510 | 1.479 | 3.58e-01 | 1.030 | 0.115 | 0.823 – 1.290 | 0.258 | 9.86e-01 |
| Neural tracking index final word [z; within-subj] | 1.032 | 0.021 | 0.991 – 1.076 | 1.517 | 3.16e-01 | 1.078 | 0.031 | 1.014 – 1.146 | 2.412 | 5.71e-02 | 0.982 | 0.028 | 0.929 – 1.038 | -0.641 | 9.38e-01 |
| ALI final word [z; within-subj] | 1.011 | 0.022 | 0.969 – 1.055 | 0.522 | 9.03e-01 | 1.022 | 0.032 | 0.960 – 1.088 | 0.678 | 7.47e-01 | 1.003 | 0.030 | 0.946 – 1.062 | 0.087 | 9.86e-01 |
| Neural tracking index final word [z; between-subj] | 1.171 | 0.067 | 1.028 – 1.335 | 2.368 | 5.62e-02 | 1.161 | 0.073 | 1.006 – 1.340 | 2.042 | 1.24e-01 | 1.185 | 0.077 | 1.019 – 1.379 | 2.197 | 1.26e-01 |
| ALI final word [z; between-subj] | 1.124 | 0.068 | 0.984 – 1.283 | 1.726 | 2.32e-01 | 1.096 | 0.074 | 0.948 – 1.268 | 1.239 | 4.30e-01 | 1.164 | 0.078 | 0.999 – 1.355 | 1.950 | 1.84e-01 |
| PTA [z] | 0.753 | 0.077 | 0.647 – 0.875 | -3.688 | **1.66e-03** | 0.760 | 0.084 | 0.645 – 0.896 | -3.273 | **4.79e-03** | 0.734 | 0.089 | 0.617 – 0.874 | -3.480 | **3.01e-03** |
| Age [z] | 0.804 | 0.079 | 0.689 – 0.938 | -2.771 | **2.46e-02** | 0.752 | 0.086 | 0.636 – 0.889 | -3.330 | **4.79e-03** | 0.868 | 0.090 | 0.727 – 1.036 | -1.565 | 3.53e-01 |
| Probed ear (right) | 1.247 | 0.067 | 1.093 – 1.422 | 3.279 | **5.72e-03** | | | | | | | | | | |
| Earlier onset (probed) | 1.000 | 0.041 | 0.923 – 1.083 | 0.005 | 9.96e-01 | 1.017 | 0.060 | 0.904 – 1.143 | 0.276 | 8.80e-01 | 0.934 | 0.057 | 0.836 – 1.044 | -1.197 | 5.20e-01 |
| Spatial cue x Semantic cue | 1.345 | 0.213 | 0.886 – 2.040 | 1.393 | 3.60e-01 | 1.355 | 0.240 | 0.847 – 2.168 | 1.269 | 4.30e-01 | 1.330 | 0.230 | 0.848 – 2.086 | 1.243 | 5.20e-01 |
| Spatial cue x ALI [within-subj] | 1.012 | 0.042 | 0.932 – 1.099 | 0.289 | 9.17e-01 | 0.950 | 0.063 | 0.840 – 1.075 | -0.809 | 7.47e-01 | 1.062 | 0.058 | 0.949 – 1.189 | 1.046 | 5.91e-01 |
| Spatial cue x Neural tracking index [within-subj] | 0.989 | 0.042 | 0.911 – 1.074 | -0.264 | 9.17e-01 | 1.003 | 0.062 | 0.888 – 1.133 | 0.049 | 9.61e-01 | 0.986 | 0.056 | 0.883 – 1.101 | -0.253 | 9.86e-01 |
| Probed ear x ALI [within-subj] | 1.038 | 0.041 | 0.957 – 1.125 | 0.897 | 6.88e-01 | | | | | | | | | | |
| Probed ear x Neural tracking index [within-subj] | 1.109 | 0.040 | 1.026 – 1.199 | 2.601 | **3.40e-02** | | | | | | | | | | |
| Earlier onset x ALI [within-subj] | 0.995 | 0.039 | 0.921 – 1.075 | -0.137 | 9.62e-01 | 1.006 | 0.058 | 0.897 – 1.127 | 0.095 | 9.61e-01 | 0.990 | 0.054 | 0.891 – 1.101 | -0.179 | 9.86e-01 |
| Earlier onset x Neural tracking index [within-subj] | 0.989 | 0.040 | 0.915 – 1.069 | -0.280 | 9.17e-01 | 0.979 | 0.058 | 0.873 – 1.098 | -0.357 | 8.66e-01 | 1.006 | 0.054 | 0.905 – 1.118 | 0.106 | 9.86e-01 |
| ALI [within-subj] x Neural tracking index [within-subj] | 1.010 | 0.021 | 0.969 – 1.053 | 0.491 | 9.03e-01 | 1.012 | 0.031 | 0.952 – 1.076 | 0.392 | 8.66e-01 | 1.005 | 0.028 | 0.951 – 1.063 | 0.183 | 9.86e-01 |
| Spatial cue x ALI [within-subj] x Neural tracking index [within-subj] | 1.019 | 0.042 | 0.938 – 1.107 | 0.444 | 9.03e-01 | 1.044 | 0.063 | 0.923 – 1.180 | 0.687 | 7.47e-01 | 1.000 | 0.057 | 0.895 – 1.118 | 0.003 | 9.97e-01 |
| Probed ear x ALI [within-subj] x Neural tracking index [within-subj] | 1.004 | 0.040 | 0.929 – 1.085 | 0.103 | 9.62e-01 | | | | | | | | | | |
| Earlier onset x ALI [within-subj] x Neural tracking index [within-subj] | 0.981 | 0.039 | 0.908 – 1.060 | -0.488 | 9.03e-01 | 0.974 | 0.059 | 0.868 – 1.093 | -0.445 | 8.66e-01 | 0.984 | 0.053 | 0.887 – 1.092 | -0.299 | 9.86e-01 |
| **Random Effects** | | | | | | | | | | | | | | | |
| $\sigma^2$ | 3.290 | | | | | 3.290 | | | | | 3.290 | | | | |
| $\tau_{00}$ | $0.552_{sentence\_pair}$ | | | | | $0.594_{sentence\_pair}$ | | | | | $0.570_{sentence\_pair}$ | | | | |
| | $0.606_{ID}$ | | | | | $0.654_{ID}$ | | | | | $0.757_{ID}$ | | | | |
| $\tau_{11}$ | $0.211_{ID.spatialSelective}$ | | | | | $0.434_{ID.spatialSelective}$ | | | | | $0.126_{ID.spatialSelective}$ | | | | |
| | $0.359_{ID.probedRight}$ | | | | | | | | | | | | | | |
| $\rho_{01}$ | $0.025_{ID.spatialSelective}$ | | | | | $0.049_{ID}$ | | | | | $-0.149_{ID}$ | | | | |
| | $-0.108_{ID.probedRight}$ | | | | | | | | | | | | | | |
| ICC | 0.283 | | | | | 0.292 | | | | | 0.292 | | | | |
| N | $155_{ID}$ | | | | | $155_{ID}$ | | | | | $155_{ID}$ | | | | |
| | $240_{sentence\_pair}$ | | | | | $240_{sentence\_pair}$ | | | | | $240_{sentence\_pair}$ | | | | |
| Observations | 35791 | | | | | 17968 | | | | | 17823 | | | | |
| Marginal $R^2$ / Conditional $R^2$ | 0.120 / 0.369 | | | | | 0.131 / 0.385 | | | | | 0.103 / 0.365 | | | | |

Note: Generalized (logistic) linear mixed-effects model, p-values for individual fixed-effect terms based on Wald test (two-sided), FDR-corrected for multiple testing.

**Supplementary Table 2: Predicting single-trial speed**

| | Speed [z] | | | | | Selective-attention trials | | | | | Divided-attention trials | | | | |
|---|---|---|---|---|---|---|---|---|---|---|---|---|---|---|---|
| *Predictors* | *Estimates* | *std. Error* | *CI* | *t-value* | *p* | *Estimates* | *std. Error* | *CI* | *t-value* | *p* | *Estimates* | *std. Error* | *CI* | *t-value* | *p* |
| Intercept | -0.037 | 0.039 | -0.113 – 0.040 | -0.938 | 4.90e-01 | 0.251 | 0.050 | 0.153 – 0.350 | 4.983 | **5.97e-06** | -0.322 | 0.035 | -0.390 – -0.253 | -9.212 | **4.79e-19** |
| Spatial cue (selective) | 0.572 | 0.039 | 0.496 – 0.648 | 14.778 | **4.49e-48** | | | | | | | | | | |
| Semantic cue (specific) | 0.198 | 0.032 | 0.136 – 0.260 | 6.279 | **2.49e-09** | 0.245 | 0.050 | 0.148 – 0.342 | 4.936 | **5.97e-06** | 0.152 | 0.039 | 0.075 – 0.228 | 3.875 | **3.99e-04** |
| Neural tracking index final word [z; within-subj] | -0.004 | 0.004 | -0.013 – 0.005 | -0.922 | 4.90e-01 | -0.012 | 0.007 | -0.024 – 0.001 | -1.774 | 1.28e-01 | 0.004 | 0.006 | -0.007 – 0.015 | 0.642 | 6.11e-01 |
| ALI final word [z; within-subj] | -0.002 | 0.004 | -0.011 – 0.007 | -0.399 | 7.59e-01 | -0.007 | 0.007 | -0.020 – 0.006 | -1.059 | 3.95e-01 | 0.004 | 0.006 | -0.008 – 0.015 | 0.629 | 6.11e-01 |
| Neural tracking index final word [z; between-subj] | 0.032 | 0.027 | -0.021 – 0.085 | 1.182 | 4.02e-01 | 0.078 | 0.044 | -0.008 – 0.165 | 1.769 | 1.28e-01 | 0.043 | 0.029 | -0.015 – 0.100 | 1.449 | 3.26e-01 |
| ALI final word [z; between-subj] | 0.006 | 0.027 | -0.047 – 0.059 | 0.211 | 8.33e-01 | 0.112 | 0.044 | 0.025 – 0.199 | 2.532 | **2.84e-02** | 0.041 | 0.029 | -0.016 – 0.099 | 1.402 | 3.26e-01 |
| PTA [z] | -0.053 | 0.031 | -0.114 – 0.008 | -1.710 | 2.13e-01 | -0.026 | 0.051 | -0.125 – 0.073 | -0.512 | 7.02e-01 | -0.046 | 0.034 | -0.111 – 0.020 | -1.360 | 3.26e-01 |
| Age [z] | -0.150 | 0.031 | -0.212 – -0.089 | -4.803 | **8.61e-06** | -0.171 | 0.051 | -0.271 – -0.071 | -3.348 | **3.06e-03** | -0.159 | 0.034 | -0.226 – -0.093 | -4.692 | **1.35e-05** |
| Probed ear (right) | 0.085 | 0.013 | 0.059 – 0.111 | 6.410 | **1.60e-09** | 0.072 | 0.017 | 0.040 – 0.105 | 4.338 | **7.20e-05** | 0.099 | 0.016 | 0.067 – 0.130 | 6.115 | **7.23e-09** |
| Earlier onset (probed) | 0.020 | 0.009 | 0.002 – 0.037 | 2.201 | 1.02e-01 | 0.018 | 0.013 | -0.007 – 0.044 | 1.405 | 2.40e-01 | 0.020 | 0.012 | -0.003 – 0.043 | 1.671 | 2.84e-01 |
| Spatial cue x Semantic cue | 0.093 | 0.063 | -0.030 – 0.217 | 1.477 | 3.07e-01 | | | | | | | | | | |
| Spatial cue x ALI [within-subj] | -0.011 | 0.009 | -0.028 – 0.006 | -1.255 | 3.95e-01 | | | | | | | | | | |
| Spatial cue x Neural tracking index [within-subj] | -0.016 | 0.009 | -0.033 – 0.001 | -1.810 | 1.93e-01 | | | | | | | | | | |
| Probed ear x ALI [within-subj] | 0.007 | 0.009 | -0.011 – 0.025 | 0.801 | 5.17e-01 | | | | | | | | | | |
| Probed ear x Neural tracking index [within-subj] | 0.009 | 0.009 | -0.008 – 0.026 | 1.022 | 4.82e-01 | | | | | | | | | | |
| Earlier onset x ALI [within-subj] | -0.017 | 0.009 | -0.034 – 0.001 | -1.896 | 1.82e-01 | -0.027 | 0.013 | -0.053 – -0.001 | -2.071 | 8.21e-02 | -0.006 | 0.011 | -0.028 – 0.017 | -0.491 | 6.68e-01 |
| Earlier onset x Neural tracking index [within-subj] | -0.003 | 0.009 | -0.020 – 0.014 | -0.343 | 7.66e-01 | 0.002 | 0.013 | -0.024 – 0.027 | 0.120 | 9.05e-01 | -0.008 | 0.011 | -0.030 – 0.014 | -0.692 | 6.11e-01 |
| ALI [within-subj] x Neural tracking index [within-subj] | -0.005 | 0.004 | -0.014 – 0.003 | -1.238 | 3.95e-01 | -0.017 | 0.007 | -0.030 – -0.004 | -2.630 | **2.56e-02** | 0.006 | 0.006 | -0.005 – 0.017 | 1.054 | 4.87e-01 |
| Spatial cue x ALI [within-subj] x Neural tracking index [within-subj] | -0.023 | 0.009 | -0.040 – -0.006 | -2.608 | **4.01e-02** | | | | | | | | | | |
| Probed ear x ALI [within-subj] x Neural tracking index [within-subj] | 0.007 | 0.009 | -0.010 – 0.024 | 0.846 | 5.15e-01 | 0.010 | 0.013 | -0.016 – 0.036 | 0.746 | 5.70e-01 | 0.004 | 0.011 | -0.018 – 0.026 | 0.343 | 7.31e-01 |
| Earlier onset x ALI [within-subj] x Neural tracking index [within-subj] | 0.005 | 0.009 | -0.012 – 0.022 | 0.592 | 6.41e-01 | 0.002 | 0.013 | -0.024 – 0.028 | 0.138 | 9.05e-01 | 0.007 | 0.011 | -0.014 – 0.029 | 0.672 | 6.11e-01 |
| **Random Effects** | | | | | | | | | | | | | | | |
| $\sigma^2$ | 0.601 | | | | | 0.710 | | | | | 0.478 | | | | |
| $\tau_{00}$ | $0.055_{sentence\_pair}$ | | | | | $0.298_{ID}$ | | | | | $0.129_{ID}$ | | | | |
| | $0.198_{ID}$ | | | | | $0.069_{sentence\_pair}$ | | | | | $0.042_{sentence\_pair}$ | | | | |
| $\tau_{11}$ | $0.077_{ID.spatialSelective}$ | | | | | $0.017_{ID.probedRight}$ | | | | | $0.020_{ID.probedRight}$ | | | | |
| | $0.015_{ID.probedRight}$ | | | | | | | | | | | | | | |
| $\rho_{01}$ | $0.717_{ID.spatialSelective}$ | | | | | $-0.136_{ID}$ | | | | | $0.114_{ID}$ | | | | |
| | $0.056_{ID.probedRight}$ | | | | | | | | | | | | | | |
| ICC | 0.319 | | | | | 0.343 | | | | | 0.270 | | | | |
| N | $155_{ID}$ | | | | | $155_{ID}$ | | | | | $155_{ID}$ | | | | |
| | $240_{sentence\_pair}$ | | | | | $120_{sentence\_pair}$ | | | | | $120_{sentence\_pair}$ | | | | |
| Observations | 32471 | | | | | 17059 | | | | | 15412 | | | | |
| Marginal $R^2$ / Conditional $R^2$ | 0.125 / 0.404 | | | | | 0.059 / 0.382 | | | | | 0.064 / 0.317 | | | | |

Note: General linear mixed-effects model, p-values for individual fixed-effect terms based on Wald test (two-sided), FDR-corrected for multiple testing.

**Supplementary Table 3: Predicting auditory alpha power lateralization during sentence presentation (3.5-6.5 s)**

| | Auditory alpha power lateralization [z] | | | | | Probed-right trials | | | | | Probed-left trials | | | | |
|---|---|---|---|---|---|---|---|---|---|---|---|---|---|---|---|
| *Predictors* | *Estimates* | *std. Error* | *CI* | *t-value* | *p* | *Estimates* | *std. Error* | *CI* | *t-value* | *p* | *Estimates* | *std. Error* | *CI* | *t-value* | *p* |
| (Intercept) | 0.000 | 0.007 | -0.013 – 0.013 | 0.054 | 9.57e-01 | -0.085 | 0.035 | -0.154 – -0.017 | -2.440 | 6.51e-02 | 0.086 | 0.037 | 0.013 – 0.159 | 2.305 | 1.91e-01 |
| Spatial cue (selective) | 0.050 | 0.009 | 0.032 – 0.069 | 5.325 | **5.54e-07** | 0.116 | 0.013 | 0.090 – 0.142 | 8.714 | **2.65e-17** | -0.016 | 0.013 | -0.042 – 0.011 | -1.166 | 5.48e-01 |
| Semantic cue (specific) | -0.010 | 0.009 | -0.028 – 0.008 | -1.058 | 5.32e-01 | -0.019 | 0.013 | -0.045 – 0.007 | -1.425 | 3.47e-01 | -0.000 | 0.013 | -0.027 – 0.026 | -0.031 | 9.75e-01 |
| Probed ear (right) | -0.171 | 0.072 | -0.312 – -0.031 | -2.390 | 6.18e-02 | | | | | | | | | | |
| Age [z] | -0.006 | 0.008 | -0.021 – 0.008 | -0.857 | 5.38e-01 | -0.041 | 0.040 | -0.121 – 0.038 | -1.023 | 4.78e-01 | 0.030 | 0.043 | -0.055 – 0.115 | 0.695 | 7.31e-01 |
| PTA [z] | 0.009 | 0.008 | -0.006 – 0.024 | 1.211 | 4.97e-01 | 0.093 | 0.041 | 0.014 – 0.172 | 2.295 | 6.51e-02 | -0.080 | 0.043 | -0.165 – 0.005 | -1.846 | 2.92e-01 |
| Earlier onset (probed) | 0.001 | 0.010 | -0.018 – 0.020 | 0.119 | 9.57e-01 | -0.009 | 0.014 | -0.036 – 0.018 | -0.667 | 5.68e-01 | 0.012 | 0.014 | -0.016 – 0.039 | 0.825 | 7.31e-01 |
| Spatial cue x Semantic cue | -0.016 | 0.019 | -0.053 – 0.021 | -0.857 | 5.38e-01 | -0.024 | 0.027 | -0.076 – 0.028 | -0.893 | 4.78e-01 | -0.008 | 0.027 | -0.061 – 0.044 | -0.317 | 8.45e-01 |
| Spatial cue x Probed ear | 0.131 | 0.019 | 0.094 – 0.168 | 6.948 | **4.08e-11** | | | | | | | | | | |
| Age x Spatial cue | -0.006 | 0.011 | -0.027 – 0.015 | -0.543 | 7.18e-01 | -0.006 | 0.015 | -0.036 – 0.024 | -0.396 | 6.92e-01 | -0.006 | 0.015 | -0.036 – 0.025 | -0.362 | 8.45e-01 |
| PTA x Spatial cue | 0.018 | 0.011 | -0.003 – 0.039 | 1.641 | 2.77e-01 | 0.014 | 0.015 | -0.016 – 0.044 | 0.934 | 4.78e-01 | 0.022 | 0.015 | -0.009 – 0.052 | 1.393 | 4.91e-01 |
| **Random Effects** | | | | | | | | | | | | | | | |
| $\sigma^2$ | 0.792 | | | | | 0.791 | | | | | 0.794 | | | | |
| $\tau_{00}$ | 0.003 $_{ID}$ | | | | | 0.183 $_{ID}$ | | | | | 0.209 $_{ID}$ | | | | |
| $\tau_{11}$ | 0.784 $_{ID.probedRight}$ | | | | | | | | | | | | | | |
| $\rho_{01}$ | -0.255 $_{ID}$ | | | | | | | | | | | | | | |
| ICC | 0.201 | | | | | 0.188 | | | | | 0.209 | | | | |
| N | 155 $_{ID}$ | | | | | 155 $_{ID}$ | | | | | 155 $_{ID}$ | | | | |
| Observations | 35791 | | | | | 17968 | | | | | 17823 | | | | |
| Marginal $R^2$ / Conditional $R^2$ | 0.009 / 0.208 | | | | | 0.010 / 0.196 | | | | | 0.005 / 0.213 | | | | |

Note: General linear mixed-effects model, p-values for individual fixed-effect terms based on Wald test (two-sided), FDR-corrected for multiple testing.

**Supplementary Table 4: Predicting neural tracking index during final word**

| | Neural tracking index [z] | | | | | Probed sentence first | | | | | Probed sentence second | | | | |
|---|---|---|---|---|---|---|---|---|---|---|---|---|---|---|---|
| *Predictors* | *Estimates* | *std. Error* | *CI* | *t-value* | *p* | *Estimates* | *std. Error* | *CI* | *t-value* | *p* | *Estimates* | *std. Error* | *CI* | *t-value* | *p* |
| (Intercept) | 0.0001 | 0.0066 | -0.0127 – 0.0130 | 0.0224 | 9.82e-01 | 0.0069 | 0.0093 | -0.0114 – 0.0252 | 0.7382 | 5.75e-01 | -0.0068 | 0.0083 | -0.0231 – 0.0095 | -0.8194 | 9.04e-01 |
| Spatial cue (selective) | 0.0691 | 0.0106 | 0.0484 – 0.0898 | 6.5451 | **7.14e-10** | 0.0426 | 0.0150 | 0.0132 – 0.0721 | 2.8368 | **4.56e-02** | 0.0955 | 0.0148 | 0.0664 – 0.1246 | 6.4373 | **1.22e-09** |
| Semantic cue (specific) | 0.0144 | 0.0106 | -0.0062 – 0.0351 | 1.3684 | 4.64e-01 | 0.0276 | 0.0150 | -0.0018 – 0.0571 | 1.8389 | 2.20e-01 | 0.0018 | 0.0148 | -0.0273 – 0.0309 | 0.1212 | 9.04e-01 |
| Probed ear (right) | -0.0305 | 0.0106 | -0.0512 – -0.0098 | -2.8880 | **2.33e-02** | -0.0297 | 0.0153 | -0.0596 – 0.0002 | -1.9447 | 2.20e-01 | -0.0340 | 0.0150 | -0.0633 – -0.0047 | -2.2737 | 1.15e-01 |
| Age (z-scored) | 0.0070 | 0.0076 | -0.0078 – 0.0219 | 0.9282 | 5.42e-01 | 0.0103 | 0.0108 | -0.0108 – 0.0314 | 0.9573 | 4.83e-01 | 0.0041 | 0.0096 | -0.0147 – 0.0228 | 0.4231 | 9.04e-01 |
| PTA (z-scored) | -0.0056 | 0.0076 | -0.0205 – 0.0093 | -0.7350 | 6.16e-01 | -0.0156 | 0.0108 | -0.0368 – 0.0055 | -1.4509 | 3.38e-01 | 0.0040 | 0.0096 | -0.0148 – 0.0228 | 0.4153 | 9.04e-01 |
| Earlier onset (probed) | 0.0137 | 0.0106 | -0.0070 – 0.0345 | 1.3013 | 4.64e-01 | | | | | | | | | | |
| Spatial cue x Semantic cue | 0.0193 | 0.0211 | -0.0221 – 0.0606 | 0.9127 | 5.42e-01 | 0.0036 | 0.0301 | -0.0554 – 0.0625 | 0.1193 | 9.05e-01 | 0.0335 | 0.0297 | -0.0247 – 0.0917 | 1.1268 | 8.66e-01 |
| Spatial cue x Probed ear | 0.0220 | 0.0211 | -0.0194 – 0.0634 | 1.0425 | 5.42e-01 | 0.0405 | 0.0301 | -0.0184 – 0.0994 | 1.3473 | 3.38e-01 | 0.0037 | 0.0297 | -0.0545 – 0.0619 | 0.1239 | 9.04e-01 |
| Age x Spatial cue | -0.0068 | 0.0122 | -0.0307 – 0.0171 | -0.5555 | 6.31e-01 | -0.0221 | 0.0174 | -0.0562 – 0.0119 | -1.2729 | 3.38e-01 | 0.0085 | 0.0171 | -0.0251 – 0.0421 | 0.4974 | 9.04e-01 |
| PTA x Spatial cue | 0.0073 | 0.0122 | -0.0166 – 0.0312 | 0.5983 | 6.31e-01 | 0.0109 | 0.0174 | -0.0232 – 0.0449 | 0.6254 | 5.91e-01 | 0.0042 | 0.0171 | -0.0294 – 0.0378 | 0.2448 | 9.04e-01 |
| Spatial cue x Earlier onset (probed) | -0.0529 | 0.0211 | -0.0943 – -0.0115 | -2.5021 | **4.94e-02** | | | | | | | | | | |
| **Random Effects** | | | | | | | | | | | | | | | |
| $\sigma^2$ | 0.9962 | | | | | 0.9895 | | | | | 1.0007 | | | | |
| $\tau_{00}$ | 0.0024 $_{ID}$ | | | | | 0.0047 $_{ID}$ | | | | | 0.0021 $_{ID}$ | | | | |
| ICC | 0.0024 | | | | | 0.0047 | | | | | 0.0021 | | | | |
| N | 155 $_{ID}$ | | | | | 155 $_{ID}$ | | | | | 155 $_{ID}$ | | | | |
| Observations | 35791 | | | | | 17588 | | | | | 18203 | | | | |
| Marginal $R^2$ / Conditional $R^2$ | 0.002 / 0.004 | | | | | 0.001 / 0.006 | | | | | 0.003 / 0.005 | | | | |

Note: General linear mixed-effects model, p-values for individual fixed-effect terms based on Wald test (two-sided), FDR-corrected for multiple testing.

### Supplementary Table 5: Predicting neural tracking index during final word

| | Neural tracking index final word [z] | | | | |
|---|---|---|---|---|---|
| *Predictors* | *Estimates* | *std. Error* | *CI* | *t-value* | *p* |
| (Intercept) | -0.0003 | 0.0066 | -0.0132 – 0.0126 | -0.0420 | 9.66e-01 |
| Auditory ALI final word [z; between-subj] | -0.0007 | 0.0065 | -0.0136 – 0.0121 | -0.1102 | 9.66e-01 |
| Auditory ALI final word [z; within-subj] | -0.0077 | 0.0054 | -0.0184 – 0.0029 | -1.4214 | 3.49e-01 |
| Spatial cue (selective attention) | 0.0709 | 0.0105 | 0.0503 – 0.0916 | 6.7236 | **1.60e-10** |
| Probed ear (right) | -0.0303 | 0.0160 | -0.0618 – 0.0012 | -1.8881 | 2.66e-01 |
| Spatial cue x ALI within-subj | -0.0165 | 0.0106 | -0.0372 – 0.0042 | -1.5596 | 3.49e-01 |
| Probed ear x ALI within-subj | -0.0059 | 0.0107 | -0.0270 – 0.0152 | -0.5490 | 7.50e-01 |
| Spatial cue x Probed ear | 0.0195 | 0.0211 | -0.0218 – 0.0609 | 0.9257 | 6.38e-01 |
| Spatial cue x Probed ear x ALI within-subj | 0.0134 | 0.0211 | -0.0281 – 0.0548 | 0.6315 | 7.50e-01 |
| **Random Effects** | | | | | |
| $\sigma^2$ | 0.9906 | | | | |
| $\tau_{00\ ID}$ | 0.0024 | | | | |
| $\tau_{11\ ID.probedRight}$ | 0.0226 | | | | |
| $\rho_{01\ ID}$ | -0.2386 | | | | |
| ICC | 0.0081 | | | | |
| $N_{ID}$ | 155 | | | | |
| Observations | 35791 | | | | |
| Marginal $R^2$ / Conditional $R^2$ | 0.002 / 0.010 | | | | |

Note: General linear mixed-effects model, p-values for individual fixed-effect terms based on Wald test (two-sided), FDR-corrected for multiple testing.

### Supplementary Table 6: Predicting auditory alpha power lateralization during final word

| | Auditory alpha power lateralization [z] | | | | | Probed-right trials | | | | | Probed-left trials | | | | |
|---|---|---|---|---|---|---|---|---|---|---|---|---|---|---|---|
| *Predictors* | *Estimates* | *std. Error* | *CI* | *t-value* | *p* | *Estimates* | *std. Error* | *CI* | *t-value* | *p* | *Estimates* | *std. Error* | *CI* | *t-value* | *p* |
| (Intercept) | 0.0003 | 0.0059 | -0.0112 – 0.0119 | 0.0589 | 9.53e-01 | -0.0590 | 0.0248 | -0.1077 – -0.0103 | -2.3749 | **4.38e-02** | 0.0594 | 0.0266 | 0.0073 – 0.1115 | 2.2337 | 1.63e-01 |
| Spatial cue (selective) | 0.0447 | 0.0100 | 0.0251 – 0.0644 | 4.4601 | **4.51e-05** | 0.1028 | 0.0142 | 0.0751 – 0.1306 | 7.2602 | **3.48e-12** | -0.0134 | 0.0142 | -0.0412 – 0.0145 | -0.9408 | 5.81e-01 |
| Semantic cue (specific) | -0.0293 | 0.0100 | -0.0490 – -0.0097 | -2.9236 | **1.27e-02** | -0.0364 | 0.0142 | -0.0642 – -0.0086 | -2.5702 | **4.38e-02** | -0.0219 | 0.0142 | -0.0497 – 0.0059 | -1.5433 | 2.76e-01 |
| Probed ear (right) | -0.1185 | 0.0505 | -0.2175 – -0.0195 | -2.3464 | 5.21e-02 | | | | | | | | | | |
| Age [z] | -0.0050 | 0.0067 | -0.0181 – 0.0082 | -0.7369 | 7.25e-01 | -0.0316 | 0.0287 | -0.0878 – 0.0246 | -1.1014 | 4.87e-01 | 0.0231 | 0.0307 | -0.0371 – 0.0832 | 0.7517 | 5.81e-01 |
| PTA [z] | 0.0066 | 0.0067 | -0.0066 – 0.0198 | 0.9802 | 5.99e-01 | 0.0672 | 0.0287 | 0.0108 – 0.1235 | 2.3367 | **4.38e-02** | -0.0580 | 0.0308 | -0.1183 – 0.0023 | -1.8857 | 1.78e-01 |
| Earlier onset (probed) | 0.0064 | 0.0105 | -0.0141 – 0.0269 | 0.6090 | 7.46e-01 | 0.0056 | 0.0148 | -0.0233 – 0.0345 | 0.3787 | 7.61e-01 | 0.0065 | 0.0148 | -0.0225 – 0.0356 | 0.4417 | 7.41e-01 |
| Spatial cue x Semantic cue | 0.0029 | 0.0200 | -0.0364 – 0.0421 | 0.1423 | 9.53e-01 | 0.0086 | 0.0283 | -0.0469 – 0.0641 | 0.3040 | 7.61e-01 | -0.0033 | 0.0284 | -0.0590 – 0.0523 | -0.1175 | 9.06e-01 |
| Spatial cue x Probed ear | 0.1162 | 0.0200 | 0.0769 – 0.1555 | 5.7970 | **7.43e-08** | | | | | | | | | | |
| Age x Spatial cue | -0.0043 | 0.0116 | -0.0270 – 0.0184 | -0.3724 | 8.67e-01 | 0.0052 | 0.0164 | -0.0269 – 0.0373 | 0.3190 | 7.61e-01 | -0.0140 | 0.0164 | -0.0461 – 0.0182 | -0.8520 | 5.81e-01 |
| PTA x Spatial cue | 0.0222 | 0.0116 | -0.0005 – 0.0450 | 1.9189 | 1.21e-01 | 0.0104 | 0.0164 | -0.0217 – 0.0424 | 0.6330 | 7.61e-01 | 0.0344 | 0.0164 | 0.0022 – 0.0666 | 2.0935 | 1.63e-01 |
| **Random Effects** | | | | | | | | | | | | | | | |
| $\sigma^2$ | 0.899 | | | | | 0.900 | | | | | 0.898 | | | | |
| $\tau_{00}$ | 0.001 $_{ID}$ | | | | | 0.088 $_{ID}$ | | | | | 0.102 $_{ID}$ | | | | |
| $\tau_{11}$ | 0.380 $_{ID.probedRight}$ | | | | | | | | | | | | | | |
| $\rho_{01}$ | -0.294 $_{ID}$ | | | | | | | | | | | | | | |
| ICC | 0.097 | | | | | 0.089 | | | | | 0.102 | | | | |
| N | 155 $_{ID}$ | | | | | 155 $_{ID}$ | | | | | 155 $_{ID}$ | | | | |
| Observations | 35791 | | | | | 17968 | | | | | 17823 | | | | |
| Marginal $R^2$ / Conditional $R^2$ | 0.005 / 0.101 | | | | | 0.006 / 0.095 | | | | | 0.003 / 0.104 | | | | |

Note: General linear mixed-effects model, p-values for individual fixed-effect terms based on Wald test (two-sided), FDR-corrected for multiple testing.

**Supplementary Table 7: Predicting neural tracking index during sentence presentation**

| | Neural tracking index [z] | | | | |
|---|---|---|---|---|---|
| *Predictors* | *Estimates* | *std. Error* | *CI* | *t-value* | *p* |
| (Intercept) | 0.0006 | 0.0064 | -0.0120 – 0.0131 | 0.0874 | 9.30e-01 |
| Spatial cue (selective) | 0.0600 | 0.0118 | 0.0368 – 0.0832 | 5.0660 | **2.24e-06** |
| Semantic cue (specific) | 0.0145 | 0.0104 | -0.0060 – 0.0349 | 1.3863 | 3.92e-01 |
| Probed ear (right) | 0.0027 | 0.0235 | -0.0435 – 0.0489 | 0.1147 | 9.30e-01 |
| Age (z-scored) | 0.0092 | 0.0074 | -0.0053 – 0.0237 | 1.2435 | 3.92e-01 |
| PTA (z-scored) | -0.0098 | 0.0074 | -0.0243 – 0.0048 | -1.3164 | 3.92e-01 |
| Earlier onset (probed) | 0.1288 | 0.0108 | 0.1077 – 0.1500 | 11.9208 | **1.01e-31** |
| Spatial cue x semantic cue | 0.0109 | 0.0209 | -0.0300 – 0.0518 | 0.5229 | 8.26e-01 |
| Spatial cue x Probed ear | 0.0486 | 0.0209 | 0.0077 – 0.0895 | 2.3313 | 7.24e-02 |
| Age x Spatial cue | 0.0090 | 0.0136 | -0.0177 – 0.0358 | 0.6631 | 7.97e-01 |
| PTA x Spatial cue | 0.0014 | 0.0137 | -0.0253 – 0.0282 | 0.1040 | 9.30e-01 |
| **Random Effects** | | | | | |
| $\sigma^2$ | 0.9729 | | | | |
| $\tau_{00\ ID}$ | 0.0022 | | | | |
| $\tau_{11\ ID.spatialSelective}$ | 0.0049 | | | | |
| $\tau_{11\ ID.probedRight}$ | 0.0690 | | | | |
| $\rho_{01}$ | 0.6279 | | | | |
| | -0.0927 | | | | |
| ICC | 0.0208 | | | | |
| $N_{ID}$ | 155 | | | | |
| Observations | 35791 | | | | |
| Marginal $R^2$ / Conditional $R^2$ | 0.005 / 0.026 | | | | |

Note: General linear mixed-effects model, p-values for individual fixed-effect terms based on Wald test (two-sided), FDR-corrected for multiple testing.

**Supplementary Table 8: Predicting neural tracking index from ALI during spatial-cue period**

| | Neural tracking index final word [z] | | | | | Neural tracking index sentence [z] | | | | |
|---|---|---|---|---|---|---|---|---|---|---|
| *Predictors* | *Estimates* | *std. Error* | *CI* | *t-value* | *p* | *Estimates* | *std. Error* | *CI* | *t-value* | *p* |
| (Intercept) | -0.0002 | 0.0066 | -0.0130 – 0.0127 | -0.0235 | 9.81e-01 | -0.0006 | 0.0065 | -0.0132 – 0.0121 | -0.0865 | 9.31e-01 |
| Auditory ALI cue [z; between-subj] | 0.0049 | 0.0066 | -0.0079 – 0.0178 | 0.7544 | 6.24e-01 | 0.0086 | 0.0063 | -0.0038 – 0.0211 | 1.3577 | 3.93e-01 |
| Auditory ALI cue [z; within-subj] | -0.0037 | 0.0053 | -0.0141 – 0.0067 | -0.6972 | 6.24e-01 | 0.0087 | 0.0053 | -0.0017 – 0.0191 | 1.6466 | 2.99e-01 |
| Spatial cue (selective attention) | 0.0685 | 0.0106 | 0.0477 – 0.0892 | 6.4746 | **8.55e-10** | 0.0611 | 0.0119 | 0.0378 – 0.0844 | 5.1461 | **2.39e-06** |
| Probed ear (right) | -0.0300 | 0.0106 | -0.0508 – -0.0093 | -2.8400 | **2.03e-02** | 0.0062 | 0.0106 | -0.0145 – 0.0269 | 0.5882 | 7.15e-01 |
| Spatial cue x ALI within-subj | -0.0122 | 0.0106 | -0.0329 – 0.0086 | -1.1492 | 5.64e-01 | -0.0101 | 0.0106 | -0.0308 – 0.0106 | -0.9544 | 5.10e-01 |
| Probed ear x ALI within-subj | -0.0010 | 0.0107 | -0.0219 – 0.0199 | -0.0908 | 9.81e-01 | -0.0121 | 0.0107 | -0.0330 – 0.0088 | -1.1337 | 4.62e-01 |
| Spatial cue x Probed ear | 0.0186 | 0.0212 | -0.0228 – 0.0601 | 0.8803 | 6.24e-01 | 0.0463 | 0.0211 | 0.0048 – 0.0877 | 2.1888 | 1.29e-01 |
| Spatial cue x Probed ear x ALI within-subj | -0.0463 | 0.0212 | -0.0878 – -0.0048 | -2.1851 | 8.66e-02 | -0.0090 | 0.0213 | -0.0506 – 0.0327 | -0.4219 | 7.57e-01 |
| **Random Effects** | | | | | | | | | | |
| $\sigma^2$ | 0.9962 | | | | | 0.9957 | | | | |
| $\tau_{00}$ | 0.0023 $_{ID}$ | | | | | 0.0021 $_{ID}$ | | | | |
| $\tau_{11}$ | | | | | | 0.0045 $_{ID.spatialSelective}$ | | | | |
| $\rho_{01}$ | | | | | | 0.6773 $_{ID}$ | | | | |
| ICC | 0.0023 | | | | | 0.0033 | | | | |
| N | 155 $_{ID}$ | | | | | 155 $_{ID}$ | | | | |
| Observations | 35791 | | | | | 35791 | | | | |
| Marginal $R^2$ / Conditional $R^2$ | 0.002 / 0.004 | | | | | 0.001 / 0.005 | | | | |

Note: General linear mixed-effects model, p-values for individual fixed-effect terms based on Wald test (two-sided), FDR-corrected for multiple testing.

**Supplementary Table 9: Predicting neural tracking index during sentence presentation**

| | Neural tracking index [z] | | | | |
|---|---|---|---|---|---|
| *Predictors* | *Estimates* | *std. Error* | *CI* | *t-value* | *p* |
| (Intercept) | -0.0001 | 0.0065 | -0.0128 – 0.0126 | -0.0124 | 9.90e-01 |
| Auditory ALI sentence [z; between-subj] | 0.0058 | 0.0064 | -0.0068 – 0.0183 | 0.9040 | 5.49e-01 |
| Auditory ALI sentence [z; within-subj] | 0.0074 | 0.0053 | -0.0030 – 0.0178 | 1.3906 | 2.96e-01 |
| Spatial cue (selective attention) | 0.0631 | 0.0119 | 0.0397 – 0.0864 | 5.2844 | **1.13e-06** |
| Probed ear (right) | 0.0067 | 0.0106 | -0.0141 – 0.0275 | 0.6319 | 6.78e-01 |
| Spatial cue x ALI within-subj | -0.0246 | 0.0106 | -0.0454 – -0.0038 | -2.3159 | 9.25e-02 |
| Probed ear x ALI within-subj | 0.0045 | 0.0110 | -0.0170 – 0.0260 | 0.4085 | 7.68e-01 |
| Spatial cue x Probed ear | 0.0414 | 0.0212 | -0.0001 – 0.0829 | 1.9531 | 1.49e-01 |
| Spatial cue x Probed ear x ALI within-subj | 0.0398 | 0.0216 | -0.0026 – 0.0822 | 1.8382 | 1.49e-01 |
| **Random Effects** | | | | | |
| $\sigma^2$ | 0.9955 | | | | |
| $\tau_{00\ ID}$ | 0.0022 | | | | |
| $\tau_{11\ ID.spatialSelective}$ | 0.0046 | | | | |
| $\rho_{01\ ID}$ | 0.6155 | | | | |
| ICC | 0.0033 | | | | |
| $N_{ID}$ | 155 | | | | |
| Observations | 35791 | | | | |
| Marginal $R^2$ / Conditional $R^2$ | 0.001 / 0.005 | | | | |

Note: General linear mixed-effects model, p-values for individual fixed-effect terms based on Wald test (two-sided), FDR-corrected for multiple testing.

**Supplementary Table 10: Predicting neural tracking index during final word**

| | Neural tracking index [z] | | | | |
|---|---|---|---|---|---|
| *Predictors* | *Estimates* | *std. Error* | *CI* | *t-value* | *p* |
| (Intercept) | 0.0001 | 0.0066 | -0.0128 – 0.0130 | 0.0219 | 9.83e-01 |
| Inferior parietal ALI final word [z; between-subj] | -0.0020 | 0.0065 | -0.0148 – 0.0109 | -0.3017 | 8.58e-01 |
| Inferior parietal ALI final word [z; within-subj] | -0.0080 | 0.0055 | -0.0189 – 0.0028 | -1.4467 | 4.44e-01 |
| Spatial cue (selective attention) | 0.0696 | 0.0105 | 0.0489 – 0.0903 | 6.5981 | **3.75e-10** |
| Probed ear (right) | -0.0307 | 0.0160 | -0.0621 – 0.0006 | -1.9208 | 2.46e-01 |
| Spatial cue x ALI within-subj | 0.0044 | 0.0106 | -0.0164 – 0.0251 | 0.4133 | 8.58e-01 |
| Probed ear x ALI within-subj | 0.0095 | 0.0109 | -0.0118 – 0.0308 | 0.8746 | 6.87e-01 |
| Spatial cue x Probed ear | 0.0212 | 0.0211 | -0.0202 – 0.0625 | 1.0029 | 6.87e-01 |
| Spatial cue x Probed ear x ALI within-subj | -0.0146 | 0.0211 | -0.0560 – 0.0269 | -0.6889 | 7.36e-01 |
| **Random Effects** | | | | | |
| $\sigma^2$ | 0.9907 | | | | |
| $\tau_{00\ ID}$ | 0.0024 | | | | |
| $\tau_{11\ ID.probedRight}$ | 0.0224 | | | | |
| $\rho_{01\ ID}$ | -0.2405 | | | | |
| ICC | 0.0080 | | | | |
| $N_{ID}$ | 155 | | | | |
| Observations | 35791 | | | | |
| Marginal $R^2$ / Conditional $R^2$ | 0.002 / 0.010 | | | | |

Note: General linear mixed-effects model, p-values for individual fixed-effect terms based on Wald test (two-sided), FDR-corrected for multiple testing.

**Supplementary Table 11: Predicting neural tracking index during sentence presentation**

| | Neural tracking index [z] | | | | |
|---|---|---|---|---|---|
| *Predictors* | *Estimates* | *std. Error* | *CI* | *t-value* | *p* |
| (Intercept) | -0.0000 | 0.0065 | -0.0127 – 0.0127 | -0.0025 | 9.98e-01 |
| Inferior parietal ALI sentence [z; between-subj] | -0.0057 | 0.0065 | -0.0183 – 0.0070 | -0.8748 | 7.72e-01 |
| Inferior parietal ALI sentence [z; within-subj] | -0.0042 | 0.0053 | -0.0146 – 0.0062 | -0.7907 | 7.72e-01 |
| Spatial cue (selective attention) | 0.0622 | 0.0106 | 0.0414 – 0.0830 | 5.8628 | **4.09e-08** |
| Probed ear (right) | 0.0045 | 0.0106 | -0.0163 – 0.0253 | 0.4271 | 8.63e-01 |
| Spatial cue x ALI within-subj | -0.0130 | 0.0106 | -0.0338 – 0.0078 | -1.2245 | 6.62e-01 |
| Probed ear x ALI within-subj | 0.0033 | 0.0113 | -0.0188 – 0.0255 | 0.2967 | 8.63e-01 |
| Spatial cue x Probed ear | 0.0446 | 0.0212 | 0.0030 – 0.0862 | 2.1026 | 1.60e-01 |
| Spatial cue x Probed ear x ALI within-subj | 0.0064 | 0.0212 | -0.0352 – 0.0481 | 0.3020 | 8.63e-01 |
| **Random Effects** | | | | | |
| $\sigma^2$ | 0.9969 | | | | |
| $\tau_{00\ ID}$ | 0.0022 | | | | |
| ICC | 0.0022 | | | | |
| $N_{ID}$ | 155 | | | | |
| Observations | 35791 | | | | |
| Marginal $R^2$ / Conditional $R^2$ | 0.001 / 0.003 | | | | |

Note: General linear mixed-effects model, p-values for individual fixed-effect terms based on Wald test (two-sided), FDR-corrected for multiple testing.

**Supplementary Table 12: Predicting single-trial accuracy**

| | Accuracy | | | | |
|---|---|---|---|---|---|
| *Predictors* | *Odds Ratios* | *std. Error* | *CI* | *z-value* | *p* |
| Intercept | 19.178 | 0.085 | 16.237 – 22.651 | 34.780 | **1.07e-263** |
| Spatial cue (selective) | 3.410 | 0.117 | 2.711 – 4.289 | 10.477 | **1.21e-24** |
| Semantic cue (specific) | 1.099 | 0.106 | 0.892 – 1.353 | 0.885 | 5.52e-01 |
| Neural tracking index sentence [z; within-subj] | 1.058 | 0.021 | 1.015 – 1.103 | 2.659 | **2.87e-02** |
| ALI sentence [z; within-subj] | 1.000 | 0.023 | 0.957 – 1.046 | 0.022 | 9.83e-01 |
| Neural tracking index sentence [z; between-subj] | 1.128 | 0.068 | 0.988 – 1.288 | 1.784 | 1.82e-01 |
| ALI sentence [z; between-subj] | 1.125 | 0.068 | 0.985 – 1.284 | 1.736 | 1.82e-01 |
| PTA [z] | 0.757 | 0.078 | 0.650 – 0.882 | -3.570 | **2.62e-03** |
| Age [z] | 0.800 | 0.080 | 0.684 – 0.934 | -2.812 | **2.17e-02** |
| Probed ear (right) | 1.236 | 0.067 | 1.084 – 1.409 | 3.156 | **8.78e-03** |
| Earlier onset (probed) | 0.993 | 0.041 | 0.916 – 1.076 | -0.177 | 9.83e-01 |
| Spatial cue x Semantic cue | 1.347 | 0.213 | 0.887 – 2.043 | 1.399 | 3.24e-01 |
| Spatial cue x ALI [within-subj] | 1.002 | 0.043 | 0.921 – 1.089 | 0.041 | 9.83e-01 |
| Spatial cue x Neural tracking index [within-subj] | 1.004 | 0.042 | 0.925 – 1.091 | 0.107 | 9.83e-01 |
| Probed ear x ALI [within-subj] | 0.961 | 0.044 | 0.882 – 1.047 | -0.903 | 5.52e-01 |
| Probed ear x Neural tracking index [within-subj] | 1.090 | 0.040 | 1.007 – 1.179 | 2.143 | 8.83e-02 |
| Earlier onset x ALI [within-subj] | 1.007 | 0.040 | 0.931 – 1.089 | 0.169 | 9.83e-01 |
| Earlier onset x Neural tracking index [within-subj] | 0.950 | 0.040 | 0.878 – 1.027 | -1.290 | 3.34e-01 |
| ALI [within-subj] x Neural tracking index [within-subj] | 1.029 | 0.022 | 0.986 – 1.074 | 1.331 | 3.34e-01 |
| Spatial cue x ALI [within-subj] x Neural tracking index [within-subj] | 1.028 | 0.043 | 0.944 – 1.118 | 0.631 | 7.26e-01 |
| Probed ear x ALI [within-subj] x Neural tracking index [within-subj] | 1.002 | 0.040 | 0.926 – 1.084 | 0.039 | 9.83e-01 |
| Earlier onset x ALI [within-subj] x Neural tracking index [within-subj] | 0.910 | 0.040 | 0.841 – 0.984 | -2.359 | 5.76e-02 |
| **Random Effects** | | | | | |
| $\sigma^2$ | 3.290 | | | | |
| $\tau_{00\ sentence\_pair}$ | 0.552 | | | | |
| $\tau_{00\ ID}$ | 0.619 | | | | |
| $\tau_{11\ ID.spatialSelective}$ | 0.214 | | | | |
| $\tau_{11\ ID.probedRight}$ | 0.355 | | | | |
| $\rho_{01\ ID.spatialSelective}$ | 0.036 | | | | |
| $\rho_{01\ ID.probedRight}$ | -0.147 | | | | |
| ICC | 0.285 | | | | |
| $N_{ID}$ | 155 | | | | |
| $N_{sentence\_pair}$ | 240 | | | | |
| Observations | 35791 | | | | |
| Marginal $R^2$ / Conditional $R^2$ | 0.120 / 0.371 | | | | |

Note: Generalized (logistic) linear mixed-effects model, p-values for individual fixed-effect terms based on Wald test (two-sided), FDR-corrected for multiple testing.

**Supplementary Table 13: Predicting single-trial speed**

| | Speed [z] | | | | |
|---|---|---|---|---|---|
| *Predictors* | *Estimates* | *std. Error* | *CI* | *t-value* | *p* |
| Intercept | -0.035 | 0.038 | -0.110 – 0.040 | -0.909 | 5.33e-01 |
| Spatial cue (selective) | 0.571 | 0.039 | 0.495 – 0.647 | 14.763 | **5.54e-48** |
| Semantic cue (specific) | 0.198 | 0.032 | 0.136 – 0.260 | 6.272 | **2.62e-09** |
| Neural tracking index sentence [z; within-subj] | -0.002 | 0.004 | -0.011 – 0.007 | -0.443 | 7.45e-01 |
| ALI sentence [z; within-subj] | -0.000 | 0.004 | -0.009 – 0.008 | -0.071 | 9.43e-01 |
| Neural tracking index sentence [z; between-subj] | 0.081 | 0.026 | 0.029 – 0.132 | 3.050 | **1.01e-02** |
| ALI sentence [z; between-subj] | 0.004 | 0.026 | -0.047 – 0.056 | 0.165 | 9.10e-01 |
| PTA [z] | -0.048 | 0.030 | -0.107 – 0.011 | -1.590 | 2.54e-01 |
| Age [z] | -0.155 | 0.031 | -0.215 – -0.096 | -5.093 | **1.94e-06** |
| Probed ear (right) | 0.085 | 0.009 | 0.068 – 0.102 | 9.769 | **1.68e-21** |
| Earlier onset (probed) | 0.023 | 0.009 | 0.005 – 0.040 | 2.558 | **3.86e-02** |
| Spatial cue x Semantic cue | 0.094 | 0.063 | -0.030 – 0.217 | 1.482 | 2.77e-01 |
| Spatial cue x ALI [within-subj] | -0.008 | 0.009 | -0.025 – 0.009 | -0.946 | 5.33e-01 |
| Spatial cue x Neural tracking index [within-subj] | -0.012 | 0.009 | -0.029 – 0.005 | -1.342 | 3.29e-01 |
| Probed ear x ALI [within-subj] | 0.018 | 0.010 | -0.001 – 0.037 | 1.850 | 2.02e-01 |
| Probed ear x Neural tracking index [within-subj] | -0.004 | 0.009 | -0.021 – 0.014 | -0.416 | 7.45e-01 |
| Earlier onset x ALI [within-subj] | -0.014 | 0.009 | -0.031 – 0.003 | -1.574 | 2.54e-01 |
| Earlier onset x Neural tracking index [within-subj] | -0.004 | 0.009 | -0.021 – 0.014 | -0.423 | 7.45e-01 |
| ALI [within-subj] x Neural tracking index [within-subj] | -0.003 | 0.004 | -0.012 – 0.006 | -0.636 | 7.21e-01 |
| Spatial cue x ALI [within-subj] x Neural tracking index [within-subj] | -0.015 | 0.009 | -0.033 – 0.002 | -1.745 | 2.23e-01 |
| Probed ear x ALI [within-subj] x Neural tracking index [within-subj] | -0.010 | 0.009 | -0.028 – 0.007 | -1.177 | 4.05e-01 |
| Earlier onset x ALI [within-subj] x Neural tracking index [within-subj] | 0.004 | 0.009 | -0.013 – 0.021 | 0.455 | 7.45e-01 |
| **Random Effects** | | | | | |
| $\sigma^2$ | 0.605 | | | | |
| $\tau_{00\ sentence\_pair}$ | 0.055 | | | | |
| $\tau_{00\ ID}$ | 0.189 | | | | |
| $\tau_{11\ ID.spatialSelective}$ | 0.077 | | | | |
| $\rho_{01\ ID}$ | 0.719 | | | | |
| ICC | 0.307 | | | | |
| $N_{ID}$ | 155 | | | | |
| $N_{sentence\_pair}$ | 240 | | | | |
| Observations | 32471 | | | | |
| Marginal $R^2$ / Conditional $R^2$ | 0.130 / 0.397 | | | | |

Note: General linear mixed-effects model, p-values for individual fixed-effect terms based on Wald test (two-sided), FDR-corrected for multiple testing.

**Supplementary Table 14: Predicting single-trial accuracy including neural tracking index based on attended decoder only**

| | Accuracy | | | | |
|---|---|---|---|---|---|
| *Predictors* | *Odds Ratios* | *std. Error* | *CI* | *z-value* | *p* |
| Intercept | 19.145 | 0.085 | 16.217 – 22.602 | 34.858 | **7.01e-265** |
| Spatial cue (selective) | 3.395 | 0.117 | 2.700 – 4.267 | 10.470 | **1.31e-24** |
| Semantic cue (specific) | 1.100 | 0.106 | 0.893 – 1.354 | 0.895 | 6.43e-01 |
| Neural tracking index final word [z; within-subj] | 1.061 | 0.022 | 1.017 – 1.107 | 2.736 | **2.28e-02** |
| ALI final word [z; within-subj] | 1.013 | 0.022 | 0.971 – 1.057 | 0.589 | 8.42e-01 |
| Neural tracking index final word [z; between-subj] | 1.141 | 0.068 | 0.998 – 1.305 | 1.935 | 1.66e-01 |
| ALI final word [z; between-subj] | 1.120 | 0.068 | 0.980 – 1.279 | 1.666 | 2.63e-01 |
| PTA [z] | 0.738 | 0.078 | 0.634 – 0.859 | -3.918 | **6.55e-04** |
| Age [z] | 0.801 | 0.080 | 0.686 – 0.937 | -2.785 | **2.28e-02** |
| Probed ear (right) | 1.242 | 0.067 | 1.088 – 1.417 | 3.220 | **7.04e-03** |
| Earlier onset (probed) | 1.000 | 0.041 | 0.923 – 1.084 | 0.006 | 9.95e-01 |
| Spatial cue x Semantic cue | 1.344 | 0.212 | 0.886 – 2.038 | 1.392 | 4.00e-01 |
| Spatial cue x ALI [within-subj] | 1.012 | 0.042 | 0.932 – 1.099 | 0.278 | 9.04e-01 |
| Spatial cue x Neural tracking index [within-subj] | 1.014 | 0.043 | 0.931 – 1.103 | 0.311 | 9.04e-01 |
| Probed ear x ALI [within-subj] | 1.037 | 0.041 | 0.956 – 1.124 | 0.877 | 6.43e-01 |
| Probed ear x Neural tracking index [within-subj] | 1.045 | 0.041 | 0.965 – 1.132 | 1.076 | 6.20e-01 |
| Earlier onset x ALI [within-subj] | 0.994 | 0.040 | 0.920 – 1.074 | -0.151 | 9.54e-01 |
| Earlier onset x Neural tracking index [within-subj] | 0.985 | 0.041 | 0.909 – 1.066 | -0.385 | 9.04e-01 |
| ALI [within-subj] x Neural tracking index [within-subj] | 1.021 | 0.021 | 0.979 – 1.065 | 0.981 | 6.43e-01 |
| Spatial cue x ALI [within-subj] x Neural tracking index [within-subj] | 0.995 | 0.043 | 0.915 – 1.082 | -0.113 | 9.54e-01 |
| Probed ear x ALI [within-subj] x Neural tracking index [within-subj] | 0.978 | 0.040 | 0.903 – 1.058 | -0.562 | 8.42e-01 |
| Earlier onset x ALI [within-subj] x Neural tracking index [within-subj] | 1.016 | 0.040 | 0.939 – 1.099 | 0.401 | 9.04e-01 |
| **Random Effects** | | | | | |
| $\sigma^2$ | 3.290 | | | | |
| $\tau_{00}$ sentence_pair | 0.550 | | | | |
| $\tau_{00}$ ID | 0.615 | | | | |
| $\tau_{11}$ ID.spatialSelective | 0.210 | | | | |
| $\tau_{11}$ ID.probedRight | 0.358 | | | | |
| $\rho_{01}$ ID.spatialSelective | 0.027 | | | | |
| $\rho_{01}$ ID.probedRight | -0.124 | | | | |
| ICC | 0.284 | | | | |
| $N_{ID}$ | 155 | | | | |
| $N_{sentence\_pair}$ | 240 | | | | |
| Observations | 35791 | | | | |
| Marginal $R^2$ / Conditional $R^2$ | 0.119 / 0.369 | | | | |

Note: Generalized (logistic) linear mixed-effects model, p-values for individual fixed-effect terms based on Wald test (two-sided), FDR-corrected for multiple testing.

**Supplementary Table 15: Predicting single-trial accuracy including neural tracking index based on attended decoder only**

| | Accuracy | | | | |
|---|---|---|---|---|---|
| *Predictors* | *Odds Ratios* | *std. Error* | *CI* | *z-value* | *p* |
| Intercept | 19.158 | 0.085 | 16.223 – 22.624 | 34.802 | **4.96e-264** |
| Spatial cue (selective) | 3.424 | 0.117 | 2.723 – 4.306 | 10.523 | **7.43e-25** |
| Semantic cue (specific) | 1.099 | 0.106 | 0.892 – 1.353 | 0.885 | 5.91e-01 |
| Neural tracking index sentence [z; within-subj] | 1.052 | 0.022 | 1.008 – 1.097 | 2.327 | 7.32e-02 |
| ALI sentence [z; within-subj] | 0.997 | 0.023 | 0.954 – 1.042 | -0.127 | 9.89e-01 |
| Neural tracking index sentence [z; between-subj] | 1.149 | 0.068 | 1.005 – 1.314 | 2.030 | 1.33e-01 |
| ALI sentence [z; between-subj] | 1.119 | 0.068 | 0.980 – 1.277 | 1.656 | 2.69e-01 |
| PTA [z] | 0.744 | 0.077 | 0.640 – 0.866 | -3.825 | **9.57e-04** |
| Age [z] | 0.797 | 0.079 | 0.682 – 0.931 | -2.861 | **1.86e-02** |
| Probed ear (right) | 1.233 | 0.067 | 1.081 – 1.407 | 3.111 | **1.02e-02** |
| Earlier onset (probed) | 0.992 | 0.041 | 0.915 – 1.075 | -0.199 | 9.75e-01 |
| Spatial cue x Semantic cue | 1.342 | 0.213 | 0.885 – 2.036 | 1.386 | 4.05e-01 |
| Spatial cue x ALI [within-subj] | 1.001 | 0.043 | 0.921 – 1.088 | 0.026 | 9.99e-01 |
| Spatial cue x Neural tracking index [within-subj] | 1.022 | 0.043 | 0.939 – 1.111 | 0.504 | 8.61e-01 |
| Probed ear x ALI [within-subj] | 0.959 | 0.044 | 0.881 – 1.045 | -0.946 | 5.83e-01 |
| Probed ear x Neural tracking index [within-subj] | 1.050 | 0.040 | 0.970 – 1.135 | 1.205 | 4.57e-01 |
| Earlier onset x ALI [within-subj] | 1.012 | 0.040 | 0.936 – 1.095 | 0.307 | 9.27e-01 |
| Earlier onset x Neural tracking index [within-subj] | 0.949 | 0.040 | 0.877 – 1.026 | -1.317 | 4.13e-01 |
| ALI [within-subj] x Neural tracking index [within-subj] | 1.009 | 0.022 | 0.967 – 1.054 | 0.432 | 8.61e-01 |
| Spatial cue x ALI [within-subj] x Neural tracking index [within-subj] | 0.980 | 0.043 | 0.900 – 1.067 | -0.459 | 8.61e-01 |
| Probed ear x ALI [within-subj] x Neural tracking index [within-subj] | 0.963 | 0.040 | 0.890 – 1.042 | -0.946 | 5.83e-01 |
| Earlier onset x ALI [within-subj] x Neural tracking index [within-subj] | 1.000 | 0.040 | 0.925 – 1.082 | -0.001 | 9.99e-01 |
| **Random Effects** | | | | | |
| $\sigma^2$ | 3.290 | | | | |
| $\tau_{00\ sentence\_pair}$ | 0.551 | | | | |
| $\tau_{00\ ID}$ | 0.618 | | | | |
| $\tau_{11\ ID.spatialSelective}$ | 0.213 | | | | |
| $\tau_{11\ ID.probedRight}$ | 0.361 | | | | |
| $\rho_{01\ ID.spatialSelective}$ | 0.062 | | | | |
| $\rho_{01\ ID.probedRight}$ | -0.152 | | | | |
| ICC | 0.285 | | | | |
| $N_{ID}$ | 155 | | | | |
| $N_{sentence\_pair}$ | 240 | | | | |
| Observations | 35791 | | | | |
| Marginal $R^2$ / Conditional $R^2$ | 0.120 / 0.371 | | | | |

Note: Generalized (logistic) linear mixed-effects model, p-values for individual fixed-effect terms based on Wald test (two-sided), FDR-corrected for multiple testing.

**Supplementary Table 16: Predicting single-trial speed including neural tracking index based on attended decoder only**

| | Speed [z] | | | | |
|---|---|---|---|---|---|
| *Predictors* | *Estimates* | *std. Error* | *CI* | *t-value* | *p* |
| Intercept | -0.037 | 0.039 | -0.114 – 0.040 | -0.942 | 5.44e-01 |
| Spatial cue (selective) | 0.572 | 0.039 | 0.496 – 0.648 | 14.786 | **3.98e-48** |
| Semantic cue (specific) | 0.198 | 0.032 | 0.136 – 0.260 | 6.275 | **2.56e-09** |
| Neural tracking index final word [z; within-subj] | -0.004 | 0.004 | -0.013 – 0.004 | -0.965 | 5.44e-01 |
| ALI final word [z; within-subj] | -0.002 | 0.004 | -0.011 – 0.007 | -0.401 | 8.42e-01 |
| Neural tracking index final word [z; between-subj] | 0.020 | 0.028 | -0.034 – 0.074 | 0.719 | 6.50e-01 |
| ALI final word [z; between-subj] | 0.005 | 0.027 | -0.048 – 0.058 | 0.189 | 8.91e-01 |
| PTA [z] | -0.057 | 0.031 | -0.118 – 0.004 | -1.828 | 2.12e-01 |
| Age [z] | -0.149 | 0.031 | -0.210 – -0.087 | -4.743 | **1.16e-05** |
| Probed ear (right) | 0.085 | 0.013 | 0.059 – 0.111 | 6.383 | **1.90e-09** |
| Earlier onset (probed) | 0.020 | 0.009 | 0.002 – 0.038 | 2.224 | 1.15e-01 |
| Spatial cue x Semantic cue | 0.093 | 0.063 | -0.030 – 0.217 | 1.478 | 3.00e-01 |
| Spatial cue x ALI [within-subj] | -0.011 | 0.009 | -0.028 – 0.006 | -1.273 | 3.72e-01 |
| Spatial cue x Neural tracking index [within-subj] | -0.005 | 0.009 | -0.022 – 0.012 | -0.564 | 7.41e-01 |
| Probed ear x ALI [within-subj] | 0.007 | 0.009 | -0.011 – 0.025 | 0.785 | 6.34e-01 |
| Probed ear x Neural tracking index [within-subj] | 0.014 | 0.009 | -0.003 – 0.031 | 1.624 | 2.87e-01 |
| Earlier onset x ALI [within-subj] | -0.016 | 0.009 | -0.033 – 0.001 | -1.867 | 2.12e-01 |
| Earlier onset x Neural tracking index [within-subj] | -0.013 | 0.009 | -0.030 – 0.004 | -1.464 | 3.00e-01 |
| ALI [within-subj] x Neural tracking index [within-subj] | -0.001 | 0.004 | -0.009 – 0.008 | -0.205 | 8.91e-01 |
| Spatial cue x ALI [within-subj] x Neural tracking index [within-subj] | -0.012 | 0.009 | -0.029 – 0.005 | -1.440 | 3.00e-01 |
| Probed ear x ALI [within-subj] x Neural tracking index [within-subj] | 0.001 | 0.009 | -0.016 – 0.018 | 0.107 | 9.15e-01 |
| Earlier onset x ALI [within-subj] x Neural tracking index [within-subj] | 0.003 | 0.009 | -0.014 – 0.020 | 0.314 | 8.72e-01 |
| **Random Effects** | | | | | |
| $\sigma^2$ | 0.601 | | | | |
| $\tau_{00}$ sentence_pair | 0.055 | | | | |
| $\tau_{00}$ ID | 0.200 | | | | |
| $\tau_{11}$ ID.spatialSelective | 0.077 | | | | |
| $\tau_{11}$ ID.probedRight | 0.016 | | | | |
| $\rho_{01}$ ID.spatialSelective | 0.718 | | | | |
| $\rho_{01}$ ID.probedRight | 0.057 | | | | |
| ICC | 0.320 | | | | |
| $N_{ID}$ | 155 | | | | |
| $N_{sentence\_pair}$ | 240 | | | | |
| Observations | 32471 | | | | |
| Marginal $R^2$ / Conditional $R^2$ | 0.124 / 0.404 | | | | |

Note: General linear mixed-effects model, p-values for individual fixed-effect terms based on Wald test (two-sided), FDR-corrected for multiple testing.

**Supplementary Table 17: Predicting single-trial speed including neural tracking index based on attended decoder only**

| | Speed [z] | | | | |
|---|---|---|---|---|---|
| *Predictors* | *Estimates* | *std. Error* | *CI* | *t-value* | *p* |
| Intercept | -0.036 | 0.039 | -0.113 – 0.041 | -0.923 | 6.52e-01 |
| Spatial cue (selective) | 0.572 | 0.039 | 0.496 – 0.647 | 14.774 | **4.72e-48** |
| Semantic cue (specific) | 0.198 | 0.032 | 0.137 – 0.260 | 6.282 | **2.45e-09** |
| Neural tracking index sentence [z; within-subj] | -0.001 | 0.004 | -0.009 – 0.008 | -0.149 | 9.68e-01 |
| ALI sentence [z; within-subj] | 0.001 | 0.005 | -0.008 – 0.011 | 0.305 | 9.68e-01 |
| Neural tracking index sentence [z; between-subj] | 0.026 | 0.027 | -0.028 – 0.080 | 0.947 | 6.52e-01 |
| ALI sentence [z; between-subj] | 0.003 | 0.027 | -0.050 – 0.056 | 0.095 | 9.68e-01 |
| PTA [z] | -0.057 | 0.031 | -0.118 – 0.003 | -1.848 | 2.15e-01 |
| Age [z] | -0.149 | 0.031 | -0.210 – -0.088 | -4.764 | **1.05e-05** |
| Probed ear (right) | 0.085 | 0.013 | 0.059 – 0.111 | 6.419 | **1.51e-09** |
| Earlier onset (probed) | 0.020 | 0.009 | 0.002 – 0.038 | 2.237 | 1.11e-01 |
| Spatial cue x Semantic cue | 0.093 | 0.063 | -0.031 – 0.217 | 1.473 | 3.44e-01 |
| Spatial cue x ALI [within-subj] | -0.009 | 0.009 | -0.026 – 0.008 | -1.015 | 6.52e-01 |
| Spatial cue x Neural tracking index [within-subj] | -0.002 | 0.009 | -0.019 – 0.015 | -0.186 | 9.68e-01 |
| Probed ear x ALI [within-subj] | 0.018 | 0.010 | -0.001 – 0.037 | 1.822 | 2.15e-01 |
| Probed ear x Neural tracking index [within-subj] | -0.000 | 0.009 | -0.017 – 0.017 | -0.035 | 9.72e-01 |
| Earlier onset x ALI [within-subj] | -0.014 | 0.009 | -0.031 – 0.003 | -1.568 | 3.21e-01 |
| Earlier onset x Neural tracking index [within-subj] | -0.005 | 0.009 | -0.023 – 0.012 | -0.614 | 7.90e-01 |
| ALI [within-subj] x Neural tracking index [within-subj] | -0.002 | 0.004 | -0.010 – 0.007 | -0.374 | 9.68e-01 |
| Spatial cue x ALI [within-subj] x Neural tracking index [within-subj] | -0.006 | 0.009 | -0.023 – 0.011 | -0.651 | 7.90e-01 |
| Probed ear x ALI [within-subj] x Neural tracking index [within-subj] | -0.007 | 0.009 | -0.024 – 0.010 | -0.837 | 6.82e-01 |
| Earlier onset x ALI [within-subj] x Neural tracking index [within-subj] | 0.002 | 0.009 | -0.015 – 0.019 | 0.208 | 9.68e-01 |
| **Random Effects** | | | | | |
| $\sigma^2$ | 0.601 | | | | |
| $\tau_{00\ sentence\_pair}$ | 0.055 | | | | |
| $\tau_{00\ ID}$ | 0.199 | | | | |
| $\tau_{11\ ID.spatialSelective}$ | 0.077 | | | | |
| $\tau_{11\ ID.probedRight}$ | 0.016 | | | | |
| $\rho_{01\ ID.spatialSelective}$ | 0.719 | | | | |
| $\rho_{01\ ID.probedRight}$ | 0.055 | | | | |
| ICC | 0.320 | | | | |
| $N_{ID}$ | 155 | | | | |
| $N_{sentence\_pair}$ | 240 | | | | |
| Observations | 32471 | | | | |
| Marginal $R^2$ / Conditional $R^2$ | 0.124 / 0.404 | | | | |

Note: General linear mixed-effects model, p-values for individual fixed-effect terms based on Wald test (two-sided), FDR-corrected for multiple testing.

**Supplementary Table 18: Predicting neural tracking of attended speech**

| | Neural tracking attended final word [z] | | | | | Neural tracking attended sentence [z] | | | | |
|---|---|---|---|---|---|---|---|---|---|---|
| *Predictors* | *Estimates* | *std. Error* | *CI* | *t-value* | *p* | *Estimates* | *std. Error* | *CI* | *t-value* | *p* |
| (Intercept) | -0.0009 | 0.0109 | -0.0221 – 0.0204 | -0.0783 | 9.38e-01 | -0.0004 | 0.0171 | -0.0340 – 0.0332 | -0.0252 | 9.80e-01 |
| Spatial cue (selective) | 0.0789 | 0.0105 | 0.0583 – 0.0994 | 7.5335 | **5.43e-13** | 0.0870 | 0.0136 | 0.0603 – 0.1138 | 6.3847 | **9.45e-10** |
| Semantic cue (specific) | 0.0078 | 0.0105 | -0.0127 – 0.0283 | 0.7458 | 5.57e-01 | 0.0153 | 0.0102 | -0.0047 – 0.0352 | 1.4978 | 3.69e-01 |
| Probed ear (right) | -0.0183 | 0.0144 | -0.0465 – 0.0099 | -1.2701 | 3.63e-01 | 0.0232 | 0.0216 | -0.0190 – 0.0655 | 1.0774 | 4.42e-01 |
| Age (z-scored) | 0.0259 | 0.0125 | 0.0014 – 0.0504 | 2.0745 | 2.09e-01 | 0.0113 | 0.0197 | -0.0273 – 0.0500 | 0.5749 | 6.51e-01 |
| PTA (z-scored) | 0.0114 | 0.0125 | -0.0131 – 0.0359 | 0.9119 | 4.98e-01 | 0.0178 | 0.0198 | -0.0210 – 0.0566 | 0.8999 | 5.06e-01 |
| Earlier onset (probed) | 0.0128 | 0.0107 | -0.0081 – 0.0338 | 1.1978 | 3.63e-01 | 0.1831 | 0.0106 | 0.1624 – 0.2037 | 17.3497 | **2.18e-66** |
| Spatial cue x semantic cue | 0.0378 | 0.0209 | -0.0032 – 0.0789 | 1.8080 | 2.59e-01 | 0.0255 | 0.0204 | -0.0145 – 0.0654 | 1.2504 | 4.42e-01 |
| Spatial cue x Probed ear | 0.0300 | 0.0209 | -0.0110 – 0.0711 | 1.4354 | 3.33e-01 | 0.0397 | 0.0204 | -0.0002 – 0.0797 | 1.9503 | 1.87e-01 |
| Age x Spatial cue | 0.0187 | 0.0121 | -0.0050 – 0.0424 | 1.5458 | 3.33e-01 | 0.0173 | 0.0157 | -0.0136 – 0.0481 | 1.0967 | 4.42e-01 |
| PTA x Spatial cue | -0.0032 | 0.0121 | -0.0269 – 0.0205 | -0.2665 | 8.69e-01 | 0.0085 | 0.0158 | -0.0224 – 0.0393 | 0.5368 | 6.51e-01 |
| **Random Effects** | | | | | | | | | | |
| $\sigma^2$ | 0.9797 | | | | | 0.9285 | | | | |
| $\tau_{00}$ | 0.0140 $_{ID}$ | | | | | 0.0415 $_{ID}$ | | | | |
| $\tau_{11}$ | 0.0150 $_{ID.probedRight}$ | | | | | 0.0127 $_{ID.spatialSelective}$ | | | | |
| | | | | | | 0.0559 $_{ID.probedRight}$ | | | | |
| $\rho_{01}$ | -0.1512 $_{ID}$ | | | | | 0.3429 | | | | |
| | | | | | | -0.0941 | | | | |
| ICC | 0.0178 | | | | | 0.0594 | | | | |
| N | 155 $_{ID}$ | | | | | 155 $_{ID}$ | | | | |
| Observations | 35791 | | | | | 35791 | | | | |
| Marginal $R^2$ / Conditional $R^2$ | 0.003 / 0.021 | | | | | 0.012 / 0.070 | | | | |

Note: General linear mixed-effects model, p-values for individual fixed-effect terms based on Wald test (two-sided), FDR-corrected for multiple testing.

**Supplementary Table 19: Predicting neural tracking of ignored speech**

| | Neural tracking ignored final word [z] | | | | | Neural tracking ignored sentence [z] | | | | |
|---|---|---|---|---|---|---|---|---|---|---|
| *Predictors* | *Estimates* | *std. Error* | *CI* | *t-value* | *p* | *Estimates* | *std. Error* | *CI* | *t-value* | *p* |
| (Intercept) | -0.0007 | 0.0090 | -0.0182 – 0.0169 | -0.0735 | 9.62e-01 | -0.0023 | 0.0152 | -0.0322 – 0.0275 | -0.1538 | 9.19e-01 |
| Spatial cue (selective) | -0.0207 | 0.0114 | -0.0431 – 0.0017 | -1.8136 | 2.56e-01 | 0.0307 | 0.0126 | 0.0059 – 0.0555 | 2.4290 | 8.33e-02 |
| Semantic cue (specific) | -0.0150 | 0.0105 | -0.0356 – 0.0055 | -1.4320 | 2.79e-01 | -0.0057 | 0.0103 | -0.0259 – 0.0144 | -0.5574 | 7.75e-01 |
| Probed ear (right) | 0.0227 | 0.0142 | -0.0052 – 0.0505 | 1.5937 | 2.67e-01 | 0.0146 | 0.0206 | -0.0259 – 0.0550 | 0.7058 | 7.55e-01 |
| Age (z-scored) | 0.0160 | 0.0103 | -0.0042 – 0.0363 | 1.5496 | 2.67e-01 | -0.0018 | 0.0175 | -0.0360 – 0.0325 | -0.1016 | 9.19e-01 |
| PTA (z-scored) | 0.0188 | 0.0103 | -0.0015 – 0.0391 | 1.8175 | 2.56e-01 | 0.0254 | 0.0175 | -0.0090 – 0.0597 | 1.4489 | 4.24e-01 |
| Earlier onset (probed) | -0.0094 | 0.0107 | -0.0304 – 0.0117 | -0.8729 | 5.24e-01 | -0.1655 | 0.0106 | -0.1864 – -0.1447 | -15.5730 | **1.22e-53** |
| Spatial cue x semantic cue | 0.0166 | 0.0210 | -0.0245 – 0.0578 | 0.7920 | 5.24e-01 | -0.0098 | 0.0206 | -0.0501 – 0.0305 | -0.4762 | 7.75e-01 |
| Spatial cue x Probed ear | -0.0010 | 0.0210 | -0.0422 – 0.0402 | -0.0479 | 9.62e-01 | -0.0218 | 0.0206 | -0.0621 – 0.0185 | -1.0610 | 6.35e-01 |
| Age x Spatial cue | 0.0259 | 0.0132 | 0.0000 – 0.0518 | 1.9610 | 2.56e-01 | 0.0208 | 0.0146 | -0.0078 – 0.0494 | 1.4248 | 4.24e-01 |
| PTA x Spatial cue | -0.0135 | 0.0132 | -0.0394 – 0.0124 | -1.0218 | 4.82e-01 | -0.0132 | 0.0146 | -0.0418 – 0.0155 | -0.9009 | 6.74e-01 |
| **Random Effects** | | | | | | | | | | |
| $\sigma^2$ | 0.9866 | | | | | 0.9445 | | | | |
| $\tau_{00}$ | 0.0081 $_{ID}$ | | | | | 0.0319 $_{ID}$ | | | | |
| $\tau_{11}$ | 0.0031 $_{ID.spatialSelective}$ | | | | | 0.0084 $_{ID.spatialSelective}$ | | | | |
| | 0.0142 $_{ID.probedRight}$ | | | | | 0.0495 $_{ID.probedRight}$ | | | | |
| $\rho_{01}$ | 0.0665 | | | | | 0.1032 | | | | |
| | -0.0639 | | | | | 0.1429 | | | | |
| ICC | 0.0125 | | | | | 0.0468 | | | | |
| N | 155 $_{ID}$ | | | | | 155 $_{ID}$ | | | | |
| Observations | 35791 | | | | | 35791 | | | | |
| Marginal $R^2$ / Conditional $R^2$ | 0.001 / 0.014 | | | | | 0.008 / 0.054 | | | | |

Note: General linear mixed-effects model, p-values for individual fixed-effect terms based on Wald test (two-sided), FDR-corrected for multiple testing.

### Supplementary Methods

#### Sentence materials

Speech stimuli consisted of 240 pairs of short German declarative sentences of fixed syntactic structure. All sentences were five words long. They always began with a first name, followed by sequence of a transitive verb, a temporal adverb, a case- and gender-ambiguous numeral and finally a plural noun (e.g., "Anna zeichnete gestern drei Stühle" ; literal translation "Anna drew yesterday three chairs"). Each position could be filled with one of ten word alternatives (see supplemental information for the full list of used words). The number of syllables at each sentence position was held constant to control for overall sentence length. All sentence contexts (i.e., consisting of the first four words) were semantically non-predictive but yielded plausible combinations with each of the 120 different sentence-final nouns. The task-relevant, sentence-final nouns belonged to two overarching general semantic categories: natural and man-made. In each of the two general categories there were ten specific subcategories (e.g., pets, fruits, vegetables in the natural, or instruments, furniture, tools in the man-made category) that each consisted of six highly representative members derived from a pre-experiment questionnaire study. Each noun was used four times across the final pool of sentence pairs, but always combined with different sentence contexts. From all possible permutations, we created 240 sentence pairs that differed at every word position.

A trained female speaker of standard German recorded the individual sentences in a sound-attenuated recording chamber (sampling rate, 44kHz). Root mean square (rms) intensity (– 26 dB Full Scale, FS) was equalized across all individual sentences. When combining the sentence recordings per pair, we temporally aligned them by the onset of the two sentence-final nouns to ensure their simultaneous presentation. This, however led to slight differences in the onset on the individual sentences. Crucially, the range and average sentence onset difference was similar for trials in which the probed (to-be-attended) sentence began earlier and those in which the unprobed (to-be-ignored) sentence began earlier (probed first: range: 0–580 ms, 162.1 ms ± 124.6; unprobed first: 0–560 ms, 180.6 ms ± 127.2). Sentence presentation was masked by continuous speech-shaped noise at a signal-to-noise-ratio of 0 dB. Noise onset was presented with a 50 ms linear onset ramp and preceded sentence onset by 200 ms. Final speech stimuli had an average length of 2512 ms (range: 2183–2963 ms). All participants listened to the same 240 sentence pairs but in subject-specific randomized order. In addition, across participants we balanced the assignment of sentences to the right and left ear, respectively.

#### EEG data acquisition and preprocessing

Participants were seated comfortably in a dimly-lit, sound-attenuated recording booth where we recorded their EEG from 64 active electrodes mounted to an elastic cap (Ag/AgCl; ActiCap / ActiChamp, Brain Products, Gilching, Germany). Electrode impedances were kept below 30 kΩ. The signal was digitized at a sampling rate of 1000 Hz and referenced on-line to the left mastoid electrode (TP9, ground: AFz). Before task instruction, 5-min eyes-open and 5-min eyes-closed resting state EEG was recorded from each participant.

For subsequent off-line EEG data analyses, we used the EEGlab[1] (version 14_1_1b) and Fieldtrip toolboxes[2] (version 2017-04-28), together with customized Matlab scripts. Independent component analysis (ICA) using EEGlab's default runica algorithm was used to remove all non-brain signal components including eye blinks and lateral eye movements, muscle activity, heartbeats and single-channel noise. Prior to ICA, EEG data were re-referenced to the average of all EEG channels (average reference). Following ICA, trials during which the amplitude of any individual EEG channel exceeded a range of 200 microvolts were removed.

#### EEG source and forward model construction

For twenty-seven participants an T1-weighted structural magnetic resonance imaging (MRI) image (Siemens MAGNETOM Skyra 3T scanner; 1-mm isotropic voxel) was acquired as part of the overarching longitudinal study design. For these participants individual EEG source and forward models were created on the basis of each participant's T1-weighted MRI image. The T1 image of one female and one male participant were used as templates for the remaining participants.

First, anatomical images were resliced (256x256x256 voxels) and the CTF convention was imposed as their coordinate system. Next, the cortical surface of each individual (or template) T1 image was constructed using the FreeSurfer (v.6.0) function *recon-all*. The result was used to generate a cortical mesh in accordance with the Human Connectome Project (HCP) standard atlas template (available at https://github.com/Washington-University/HCPpipelines) using HCP Workbench (v1.5). We used FieldTrip (*ft_postfreesurferscript.sh*) to create the individual cortical mesh encompassing 4002 grid points per hemisphere. This representation provided the individual EEG source model geometry. To generate individual head and forward models, we segmented T1 images into three tissue types (skull, scalp, brain; Fieldtrip function *ft_volumesegment*).

Subsequently, the forward model (volume conduction) was estimated using the boundary element method in FieldTrip ('dipoli' implementation). Next, we optimized the fit of digitized EEG channel locations (xensor digitizer, ANT Neuro) to each individual's head model in the CTF coordinate system using rigid-body transformation and additional interactive alignment. Finally, the lead-field matrix for each channel x source pair was computed using the source and forward model per individual.

#### Beamforming

Source reconstruction (inverse solution) was achieved using a frequency-domain beamforming approach, namely partial and canonical coherence (PCC)[3]. To this end, we concatenated 5-min segments of eyes-open resting state and a random 5-min segment of task data and calculated the cross-spectral density using a broad-band filter centred at 15 Hz (bandwidth = 14 Hz). The result was then used together with the source and forward model to obtain a common spatial filter per individual in FieldTrip (regularization parameter: 5 %, dipole orientation: axis of most variance using singular value decomposition).

#### Regions of interest

We constrained the analysis of neural measures to an a priori defined, source-localized auditory region of interest (ROI) as well as one control ROI in the inferior parietal lobule. Regions of interest were constructed by selecting a bilaterally symmetric subset of functional parcels (see Supplementary Fig. 3). Following the notation used in ref.[4], the auditory ROI encompassed eight parcels per hemisphere covering the primary auditory cortex, lateral, medial, and parabelt complex, A4 and A5 complex that extended laterally into the posterior portion of the superior temporal gyrus (Brodmann area (BA) 22). It also included parts of the retroinsular and parainsular cortex (BA52) that lie at the boundary of temporal lobe and insula. The inferior parietal (IPL) ROI consisted of six parcels per hemisphere that included the angular gyrus (BA39), supramarginal gyrus (BA40), as well as the anterior lateral bank of the intraparietal sulcus.

#### Extraction of envelope onsets

From the presented speech signals, we derived a temporal representation of the acoustic onsets in the form of the onset envelope[5]. To this end, using the NSL toolbox [6], we first extracted an auditory spectrogram of the auditory stimuli (128 spectrally resolved sub-band envelopes logarithmically spaced between 90–4000 Hz), which were then summed across frequencies to yield a broad-band temporal envelope. Next, the output was down-sampled and low-pass filtered to match the specifics of the EEG. To derive the final onset envelope to be used in linear regression, we first obtained the first derivative of the envelope and set negative values to zero (half-wave rectification) to yield a temporal signal with positive-only values reflecting the acoustic onsets (see Fig. 4A and Supplementary Fig. 4).

#### Decoding accuracy

To evaluate how accurately we could decode an individual's focus of attention, we separately evaluated the performance of the reconstruction models for the attended and the ignored envelope. When the reconstruction was performed with the attended reconstruction models, the focus of attention was correctly decoded if the correlation with the attended envelope was greater than that with the ignored envelope. Conversely, for reconstructions performed with the ignore reconstruction models, a correct identification of attention required the correlation with the ignored envelope to be greater than that with the attended envelope (see Supplementary Fig. 10). To quantify whether the single-subject decoding accuracy was significantly above chance, we used a binomial test with $\alpha$=.05.

Given that our challenging listening task presented two concurrent sentences that were (i) of short duration, (ii) spoken by the same female talker, (iii) highly similar with respect to their onset envelopes, and (iv) presented against speech-shaped noise, we wished to further evaluate the potency of the employed envelope reconstruction approach under such difficult conditions. To this end, we examined the decoding accuracy across participants in selective-attention trials (see also Supplementary Fig. 4). Evaluated at the single-trial level, attention was correctly decoded if the envelope reconstructed by the attended reconstruction model was more similar to the attended envelope than to the ignored envelope ($r_{attended} > r_{ignored}$) and vice versa for the ignored reconstruction model.

Unsurprisingly, the achieved decoding accuracy was lower than that reported in speech tracking studies using much longer segments of continuous speech but still reached a mean decoding accuracy of 60% (range: 0.40–0.79) for the reconstruction model of the attended envelope and 57% (range: 0.41–0.75) for the reconstruction model of the ignored envelope. Importantly, when investigated at the single-subject level, the attended reconstruction model yielded decoding accuracies that exceeded the empirical chance level of 58% in 89 out of 155 participants. The ignored reconstruction model yielded significantly above chance performance in 66 out 155 participants. In addition to the aforementioned experimental conditions that created a particularly difficult scenario for stimulus reconstruction, it is important to bear in mind that decoding accuracy is not only influenced by specifics of the linear model itself but also by how closely participants followed the spatial-cue instructions. For these reasons, our analyses focus on the single-trial neural tracking strength as a more variable single-trial measure of how strongly the attended and ignored sentence were cortically represented in a particular trial.
